# Automated quantification of white matter hyperintensity confluence: A measure of spatial organisation beyond volume and visual rating scales

**DOI:** 10.64898/2026.07.24.740551

**Authors:** Tatjana Schmidt, Robert Salzmann, Marcella Montagnese, Dennis Chan, Jose Bernal, Malte Pfister, Philipp Arndt, Oliver Peters, Julian Hellmann-Regen, Lukas Preis, Daria Gref, Josef Priller, Eike Jakob Spruth, Maria Gemenetzi, Slawek Altenstein, Anja Schneider, Klaus Fliessbach, Okka Kimmich, Jens Wiltfang, Claudia Bartels, Björn H. Schott, Ayda Rostamzadeh, Wenzel Glanz, Enise I. Incesoy, Michaela Butryn, Katharina Buerger, Daniel Janowitz, Sophia Stoecklein, Robert Perneczky, Boris-Stephan Rauchmann, Stefan Teipel, Mihovil Mladinov, Alice Grazia, Christoph Laske, Sebastian Sodenkamp, Annika Spottke, Gabor C. Petzold, Michael Wagner, Falk Lüsebrink, Luca Kleineidam, Stefan Hetzer, Peter Dechent, Stefanie Schreiber, Emrah Düzel, Frank Jessen, Gabriel Ziegler, Benjamin R. Underwood, Timothy Rittman

**Affiliations:** Department of Clinical Neurosciences, University of Cambridge, Cambridge Biomedical Campus, Cambridge CB2 0QQ, UK; Université de Lyon, Inria, ENS de Lyon, LIP, 46 Allée d’Italie, 69364 Lyon Cedex 07, France; Institute for Quantum Information, RWTH Aachen University, Otto-Blumenthal-Strasse 20, 52074 Aachen, Germany; Department of Psychology, University of Cambridge, Downing Street, Cambridge CB2 3EB, UK; Institute of Cognitive Neuroscience, University College London, London, UK; Department of Psychiatry, University of Cambridge, Herschel Smith Building, Cambridge Biomedical Campus, Cambridge CB2 0SZ, UK; Cambridgeshire and Peterborough NHS Foundation Trust, Windsor Unit, Fulbourn Hospital, Cambridge CB21 5EF, UK; Department of Artificial Intelligence in Biomedical Engineering (AIBE), Faculty of Engineering, Friedrich-Alexander-Universität Erlangen-Nürnberg (FAU), Erlangen, Germany; Faculty of Medicine, Friedrich-Alexander-Universität Erlangen-Nürnberg (FAU), Erlangen, Germany; German Centre for Neurodegenerative Diseases (DZNE), Magdeburg, Germany; Institute of Cognitive Neurology and Dementia Research (IKND), Otto-von-Guericke Universität Magdeburg, Magdeburg, Germany; Institute for Neuroscience and Cardiovascular Research, Row Fogo Centre for Research into Ageing and the Brain, Department of Neuroimaging Sciences, The University of Edinburgh, Edinburgh, UK; Department of Neurology, University Hospital Magdeburg, Magdeburg, Germany; Charité Universitätsmedizin Berlin, Department of Psychiatry and Neurosciences, Berlin, Germany; Charité Universitätsmedizin Berlin, ECRC Experimental and Clinical Research Center, Berlin, Germany; German Centre for Neurodegenerative Diseases (DZNE), Berlin, Germany; Department of Psychiatry and Psychotherapy, Charité, Berlin, Germany; Department of Psychiatry and Psychotherapy, School of Medicine and Health, Technical University of Munich, and German Center for Mental Health (DZPG), Munich, Germany; University of Edinburgh and UK DRI, Edinburgh, UK; Department of Old Age Psychiatry and Cognitive Disorders, University Hospital Bonn, University of Bonn, Bonn, Germany; German Centre for Neurodegenerative Diseases (DZNE), Bonn, Germany; Department of Psychiatry and Psychotherapy, University Medical Center Goettingen, University of Goettingen, Goettingen, Germany; German Centre for Neurodegenerative Diseases (DZNE), Goettingen, Germany; Leibniz Institute for Neurobiology, Magdeburg, Germany; Neurosciences and Signaling Group, Institute of Biomedicine (iBiMED), Department of Medical Sciences, University of Aveiro, Aveiro, Portugal; Department of Psychiatry and Psychotherapy, University of Cologne, Medical Faculty, Cologne, Germany; Excellence Cluster on Cellular Stress Responses in Aging-Associated Diseases (CECAD), University of Cologne, Cologne, Germany; Department for Psychiatry and Psychotherapy, University Clinic Magdeburg, Magdeburg, Germany; German Centre for Neurodegenerative Diseases (DZNE), Munich, Germany; Institute for Stroke and Dementia Research (ISD), University Hospital, LMU Munich, Munich, Germany; Department of Radiology, Ludwig Maximilian University Hospital, Munich, Germany; Department of Psychiatry and Psychotherapy, University Hospital, LMU Munich, Munich, Germany; Munich Cluster for Systems Neurology (SyNergy), Munich, Germany; Ageing Epidemiology Research Unit (AGE), School of Public Health, Imperial College London, London, UK; Sheffield Institute for Translational Neuroscience (SITraN), University of Sheffield, Sheffield, UK; Department of Neuroradiology, University Hospital LMU, Munich, Germany; Department of Psychosomatic Medicine, Rostock University Medical Center, Rostock, Germany; German Centre for Neurodegenerative Diseases (DZNE), Rostock-Greifswald, Germany; Department of Psychosomatic Medicine, University Medicine Rostock, Rostock, Germany; Department of Psychiatry and Psychotherapy, University of Tübingen, Tuebingen, Germany; German Centre for Neurodegenerative Diseases (DZNE), Tuebingen, Germany; Section for Dementia Research, Department of Psychiatry and Psychotherapy, Hertie Institute for Clinical Brain Research, University of Tübingen, Tuebingen, Germany; Clinic for Parkinson’s, Sleep and Movement Disorders, Centre for Neurology, University Hospital Bonn, Bonn, Germany; Berlin Center for Advanced Neuroimaging, Charité – Universitätsmedizin Berlin, Berlin, Germany; MR-Research in Neurosciences, Department of Cognitive Neurology, University Medical Center Goettingen, Germany; Department of Vascular Neurology, University Hospital Bonn, Germany

**Keywords:** White Matter Hyperintensity, MRI, White Matter Pathology, Neurodegeneration, cerebral small vessel disease

## Abstract

White matter hyperintensities (WMH) are a highly prevalent finding on FLAIR MRI scans and a prominent feature of white matter pathology across cerebrovascular and neurodegenerative diseases. Currently, WMH are assessed with visual rating scales such as the Fazekas scale or with their volume, as calculated from automatic or manual segmentations. Both methods have limitations: Visual rating scales are rater-dependent and coarse, while WMH volume does not take the confluence of lesions into account and thus disregards their spatial organisation. As an alternative, here we propose a novel automated method for quantifying the confluence of white matter hyperintensities on a continuous standardised scale between 0 and 1. The metric is based on WMH segmentations from routine MRI and quantifies the extent to which individual WMH merge into coherent lesions, independently of total lesion volume.

We apply the method to QMIN-MC, a large UK memory clinic cohort, and show associations of the confluence metric with age, cognitive performance across domains, and Fazekas ratings. Participants with vascular and mixed dementia showed higher confluence than other diagnostic groups, whereas cognitively unimpaired participants showed lower confluence. However, confluence did not explain additional cognitive variance after accounting for log-transformed WMH volume. Findings were validated in DELCODE, an independent cohort of individuals with neurodegenerative disorders, replicating our original results. In this validation cohort, periventricular WMH confluence remained associated with cognition after adjustment for WMH volume.

These findings introduce WMH confluence as a reproducible, automated, and fine-grained measure of lesion spatial organisation. It provides complementary information about morphological WMH severity beyond volume and is an alternative to visual rating scales. Although related to WMH volume in memory-clinic populations, confluence captures clinically interpretable information and may complement existing WMH measures for improved lesion characterisation in studies of white matter disease, ageing, and cognitive impairment.

## 1 Introduction

White matter hyperintensities (WMH) of presumed vascular origin are a common finding on MRI scans and appear bright on FLAIR, T2-weighted and proton density images (Prins and Scheltens, 2015). Most commonly, WMH are a prominent feature of cerebral small vessel disease and are caused by chronic ischaemia (R. Schmidt et al., 2007, Wardlaw et al., 2015). They vary in size and location, ranging from (multi-)focal lesions to diffuse confluent lesions which can cover extensive areas (Sarbu et al., 2016; Wardlaw et al., 2013). Histopathological studies of WMH have revealed demyelination, axonal loss, edema and gliosis (Wardlaw et al., 2015; Gouw et al., 2011) as the underlying neuropathological changes. WMH therefore reflect tissue damage of varying severity and disruption to white matter tracts.

In adults, WMH are very common, found in between 39% and 96% of the otherwise healthy population (Prins and Scheltens, 2015). Their clinical impact is highly variable and, whilst not every individual with WMH will develop dementia (Zeestraten et al., 2017), WMH have been consistently linked to cognitive impairment (Debette and Markus, 2010; Prins and Scheltens, 2015; d’Arbeloff et al., 2019; Kloppenborg et al., 2014; Lohner et al., 2025), functional decline (Puzo et al., 2019; Jochems et al., 2024; Inzitari et al., 2009), dementia (Brickman et al., 2015; Vermeer et al., 2003) and stroke (Debette et al., 2010; Buyck et al., 2009) in the general population and in high-risk groups. Furthermore, identifying WMH is important in the emerging era of disease modifying treatments, since they increase the risk of Amyloid Related Inflammatory Abnormalities (ARIA) and haemorrhage in clinical trials of monoclonal antibodies for the treatment of Alzheimer’s disease (Cummings et al., 2023). Therefore, WMH are clinically highly relevant and there is a need for their accurate, automated and quantitative characterisation.

Current approaches to characterising WMH include volumetric assessment from manual or automated segmentation, and visual rating scales that take volume and spatial distribution into account (Fazekas et al., 1987; Scheltens et al., 1993; Wahlund et al., 2001)^1^. While studies employing WMH volume and visual rating scales have substantially advanced our understanding of white matter changes, both approaches have limitations. Visual rating scales are rater-dependent, limited in their reproducibility, and coarse. The most common scale, the Fazekas scale, ranges from 0 to 3 and therefore does not offer a fine-grained assessment of lesion burden. On the other hand, WMH volume is a rater-independent and continuous measure, but does not capture the range of lesion properties that reflect underlying mechanisms by which lesions contribute to cognitive and functional impairment (R. Schmidt et al., 2007). White matter is not a uniform entity that is linearly affected by lesion burden. Rather, it is organised in tracts which form complex networks (Catani and Schotten, 2012), making it important to consider the spatial organisation of lesions in addition to their volume.

One way this has been addressed in more recent studies is by moving beyond focal damage and focusing on the extent to which WMH disrupt white matter tracts and networks, using functional MRI (Leone et al., 2024; Crockett et al., 2021; Taylor et al., 2017; Lockhart et al., 2015; Vettore et al., 2021), diffusion MRI (Li et al., 2023; Taghvaei et al., 2024; D. Yang et al., 2020; Vergoossen et al., 2020; Rizvi et al., 2020; Kan et al., 2025; Busby et al., 2025) or both (Petersen et al., 2024; Jaywant et al., 2022). These advanced approaches are highly promising and have contributed to elucidating the relationship between WMH and cognitive impairment, however they require complex analysis and are therefore not scalable in clinical settings.

Here, we propose another approach to characterising the spatial organisation of WMH beyond volume which can be employed in the clinical context: an algorithm to automatically quantify WMH confluence on a continuous scale from 0 to 1 based on a WMH segmentation of an MRI scan. *Confluence* of WMH refers to the degree to which discrete WMH have merged into coherent lesions. It forms parts of commonly used visual rating scales, such as the Fazekas scale, and is intuitive for humans to recognise. In line with the interpretation of WMH as markers of white matter tract disruption rather than merely focal tissue damage, we hypothesise that more confluent lesions contribute to more significant cognitive impairment, presumably because they disrupt a broader network of interconnected brain regions than punctate lesions of comparable volume.

This is supported by converging evidence from histopathological, radiological and clinical studies indicating that confluent WM lesions differ from punctate (i.e. non-confluent) WM lesions in their underlying pathologies, progression rates, and relationship with cognitive and functional impairment (R. Schmidt, Grazer, et al., 2011; R. Schmidt, Schmidt, et al., 2011; Fazekas et al., 1993; Pantoni, 2010). Confluent lesions are ischemic in origin and progress more rapidly than punctate lesions (R. Schmidt et al., 2004). Punctate lesions are often not related to small vessel disease and are more benign in terms of progression and impact (R. Schmidt, Grazer, et al., 2011). Increasing confluence of lesions has been associated with increasing severity of WM damage (Gouw et al., 2011). It has also been suggested that not all punctate lesions are on a continuum with confluent lesions, and that instead they differ in their underlying aetiologies (Enzinger et al., 2006). In contrast to punctate WMH, confluent WMH have been associated with cognitive impairment (van der Flier et al., 2005), cognitive decline (Garde et al., 2005; R. Schmidt et al., 2005; van den Heuvel et al., 2006), and impairment in activities of daily living (Inzitari et al., 2009). Taken together, these findings show that WMH confluence is a clinically relevant concept. While confluence is implicitly taken into consideration in visual rating scales such as the Fazekas scale, there is currently no rater-independent automated method to quantify it on a continuous scale.

We propose an algorithm to automatically quantify WMH confluence. Our measure captures the degree of lesion confluence independently of total lesion volume, providing a complementary description of WMH spatial organisation. We apply our algorithm in a UK memory clinic population to investigate associations of WMH confluence with WMH volume, Fazekas scores, age, and cognitive performance, and to examine differences between diagnostic groups. We validate the findings in an independent cohort of neurodegenerative disorders from Germany.

## 2 Confluence metric

We propose a confluence operator, Conf(·), to quantify the extent to which individual WMH merge into continuous lesions. The operator evaluates all pairs of voxels in an image: pairs of voxels contribute greatly to the score if they are close together *and* both are classified as WMH. More distant WMH voxel pairs contribute only minimally.

### 2.1 Confluence in two dimensions

Let *A* ∈ [0, 1]*^X×Y^ ^×Z^* denote a 3D input image with *Z* 2D axial slices. Here, *A* may represent either a WMH probability map, where each voxel value corresponds to the probability of belonging to WMH, or a binary WMH segmentation mask, where voxel values are 1 if WMH is present and 0 otherwise. A binary mask should be preferred in order to make absolute confluence values comparable across studies.

We denote the *z*-th slice by *A*^(*z*)^ ∈ [0, 1]*^X×Y^*, *z* = 1, *…*, *Z*, with the matrix *A*^(*z*)^ represented in equation (1).

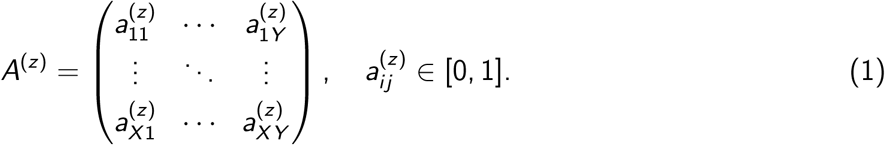

We define the operator Conf(·) as a two-body interaction over all voxel pairs in a slice, weighted by spatial distance:

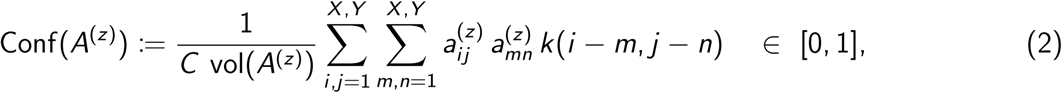

where 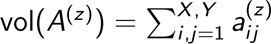 is the WMH volume in slice *z*, *C* is a normalisation constant chosen such that Conf(·) ∈ [0, 1], and *k*(·, ·) is a distance-dependent interaction kernel. Each voxel pair (*i*, *j*) and (*m*, *n*) contributes proportionally to the product of their WMH probabilities (in the case of probabilistic WMH maps) or their WMH occupancy (in the case of binary WMH maps) *a_ij_*^(*z*)^, *a_min_*^(*z*)^ ∈ [0, 1], with *_i_*contribution scaled by their distance, *k*(*i* − *m*, *j* − *n*). Thus, neighbouring WMH voxels exert stronger interactions than distant ones.

**Interaction kernel.** We employ a Gaussian interaction kernel *k*(*i* −*m*, *j* −*n*) = exp(−*s*[(*i* −*m*)^2^+(*j* −*n*)^2^]) with *s* = 0.05. This parameter value for *s* was chosen empirically so that interactions beyond a radius of 10 voxels contribute less than 0.01, which reflects a plausible spatial scale in the brain. The interaction terms 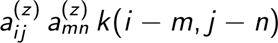 are then summed over all voxel pairs.

**Normalisation.** Without normalisation by the WMH volume of a slice, the confluence score would increase with the number of WMH voxels, irrespective of their spatial distribution. The normalisation by volume in equation (2) ensures that the measure reflects only the degree of confluence.

**Scaling.** For interpretability and comparability, the confluence metric is also scaled to the interval [0, 1] by dividing by the maximum attainable confluence metric *C*, corresponding to an image fully occupied by WMH, see equation (3).

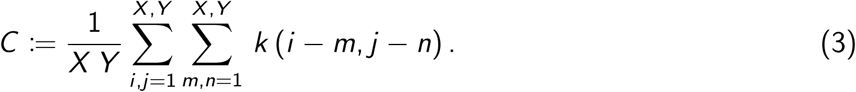

**Calculating confluence for an entire volume.** As a final step, in order to obtain a confluence metric for an entire image *A* ∈ [0, 1]*^X×Y^ ^×Z^*, the volume-normalised and scaled Conf(*A*) values of all slices in a volume are summed and divided by the number of slices containing WMH to obtain the average confluence metric for the image.

### 2.2 Confluence in three dimensions

An alternative to the 2-dimensional approach outlined above is to calculate confluence in 3D instead, using equation (4) with the interaction kernel in equation (5) and WMH volume defined in equation (6). The maximum attainable value of confluence metric C (used for scaling) is defined in equation (7).

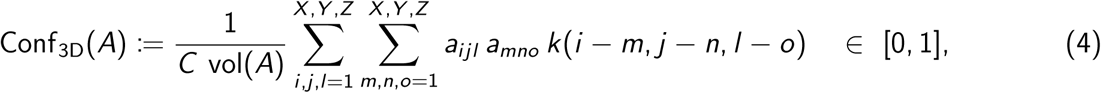

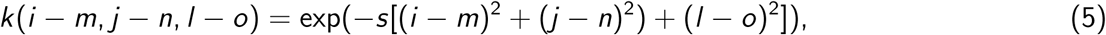

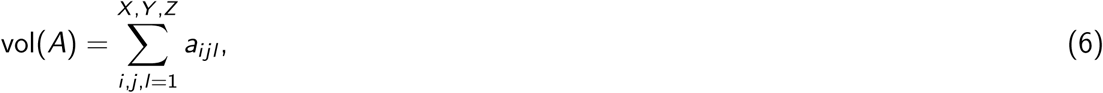

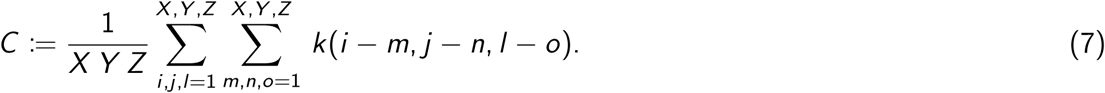

### 2.3 Demonstration of the confluence metric’s properties

Figure 1a demonstrates the metric’s properties with synthetic lesion images. The “lesion” volume of image B is twice that of image A, but the volume-normalisation ensures that the confluence metrics are nearly identical (0.14 and 0.17), reflecting that the lesions in A and B do not differ in their degree of confluence. The volume in image C is identical to that in B, but as lesions are closer to each other, C has higher confluence (0.25). Finally, D has twice the lesion volume of C, but the confluence metric of D is less than twice that of C (0.40), reflecting that the lesion topology in C and D is similar and that the degree of confluence is not quite twice as high. As lesions approach each other or start merging, the confluence metric increases consistently up to a value of 1. Figure 1b illustrates this with exemplary image slices from study participants. While image A and B have similar lesion volumes, visual inspection suggests that the spatial organisation differs between the two images and that they should obtain different scores on the Fazekas scale (grade 1 and 2, respectively). The confluence metric reflects these differences and distinguishes more clearly between the two images.

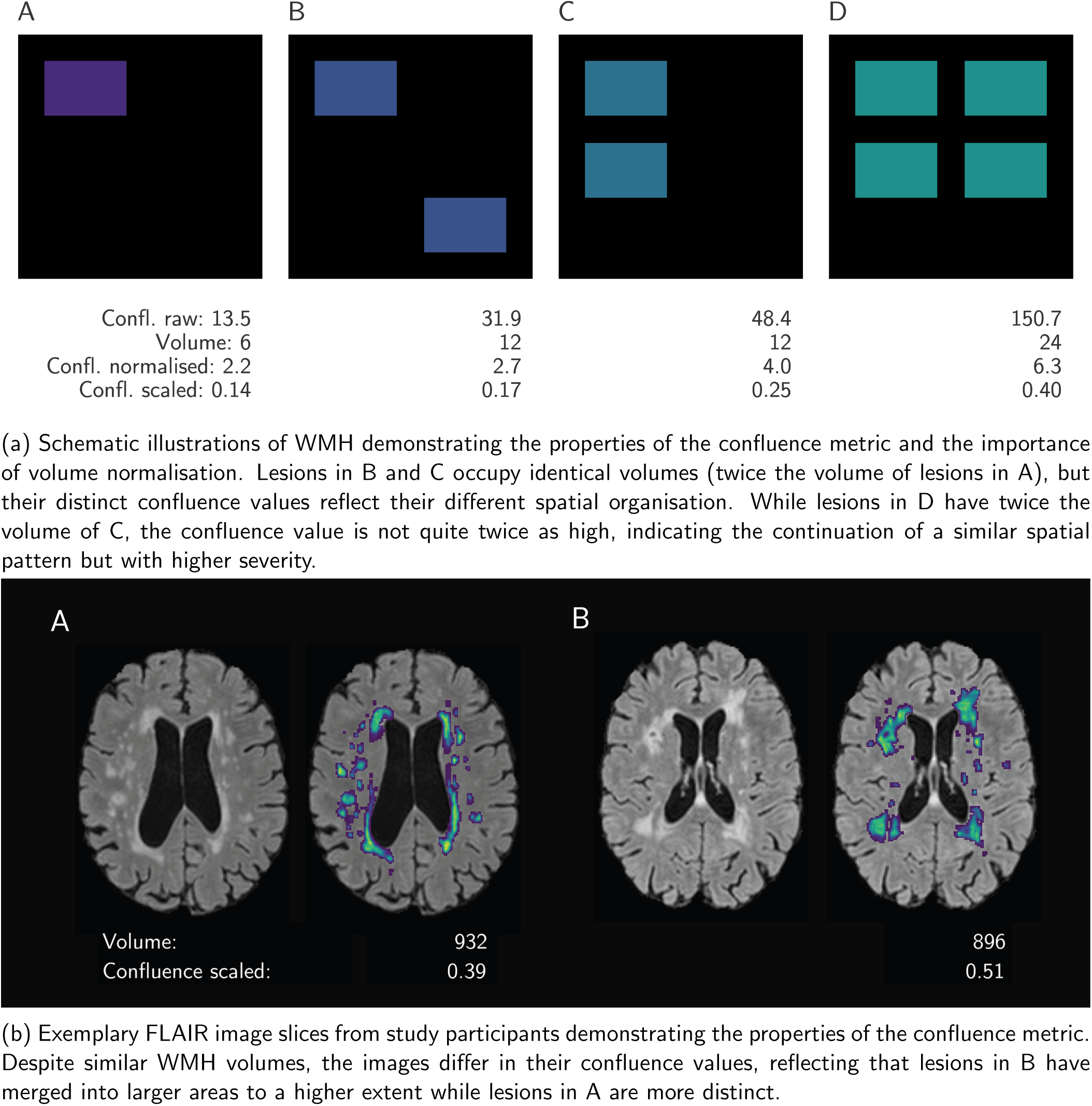

## 3 Methods

### 3.1 Participants

Participants were recruited to the QMIN-MC study from memory clinic services in two NHS Trusts (Cambridgeshire and Peterborough NHS Foundation Trust, Cambridge University Hospitals NHS Foundation Trust). All participants were eligible regardless of their age, gender or co-morbidities as long as they had no contraindications to undergoing MRI. These open inclusion criteria ensure that the cohort is as close to a real-life cohort encountered in memory clinics as possible. 1073 participants were included (537 male, 529 female, 7 other, mean age 72 ± 11 years, age range 26–96 years). Written informed consent was obtained and the study was approved by the East of England - Essex Research Ethics Committee.

### 3.2 Cognitive assessment

Clinical data were collected from all participants and included age, clinical diagnosis, years of education and scores from cognitive tests (Addenbrooke’s Cognitive Examination Revised and Addenbrooke’s Cognitive Examination III, which assess the five cognitive domains attention, memory, language, verbal fluency and visuospatial function; Mioshi et al., 2006; Noone, 2015). Different but equivalent ACE versions were administered at different study sites (ACE-R or ACE III); they were analysed together and will be referred to as “ACE”.

### 3.3 Diagnosis

The dataset contains participants with a range of diagnoses. Participants were diagnosed by consultants with a special interest in dementia as part of their memory clinic assessment according to international diagnostic guidelines, and diagnoses were confirmed at multidisciplinary team meetings. Diagnoses were further grouped before entering analysis because some diagnoses were shared by less than ten participants. The following diagnostic groups were used:

- Alzheimer’s disease (AD, including posterior cortical atrophy and logopenic aphasia, n = 283),
- non-AD dementia (including frontotemporal dementia, dementia with Lewy bodies, Parkinson’s disease dementia, progressive supranuclear palsy, corticobasal syndrome; n = 28),
- vascular dementia (n = 29),
- mixed dementia (AD with vascular dementia; n = 53),
- mild cognitive impairment (MCI, n = 141),
- other diagnoses (psychiatric disorders, motor disorder without dementia, other cognitive disorders, uncertain/unknown diagnosis; n = 362),
- no cognitive impairment (functional/attentional memory symptoms, healthy; n = 177),

### 3.4 MRI acquisition

MR images were acquired on a 3T whole-body MRI system (Magnetom Prisma, Siemens Medical Systems, Germany) using an 80 mT/m gradient coil and a 64-channel radio-frequency receive head coil at the Wolfson Brain Imaging Centre, Cambridge. 3D MP-RAGE images (Magnetisation-Prepared RApid Gradient Echo) were acquired with isotropic voxel size = 1.0 mm^3^, TR = 2000 ms, TE = 1.95 ms, TI = 880 ms, flip angle = 8°, FOV = 256 × 256 × 208 mm, 208 sagittal slices, GRAPPA reconstruction and acceleration factor = 2. FLAIR images (FLuid-Attenuated Inversion Recovery) were acquired with voxel size 1.0 × 1.0 × 1.05 mm, TR = 5000 ms, TE = 386 ms, TI = 1800 ms, FOV = 256 × 256 × 202 mm, 192 sagittal slices, GRAPPA reconstruction and acceleration factor = 3, partial Fourier factor = 7/8. The total acquisition time was 9:40 minutes.

### 3.5 MRI preprocessing, WMH segmentation and WMH confluence

T1-weighted and FLAIR images were preprocessed with micapipe version 0.2.3 (Cruces et al., 2022) using default settings. Total intracranial volume was derived with FreeSurfer (Fischl, 2012 using the procedure from Buckner et al. (2004). WMH were segmented using T1-weighted and FLAIR images with FSL’s TrUE-Net toolbox and its pretrained model (Triplanar U-Net ensemble network model; Sundaresan, Zamboni, Rothwell, et al., 2021; Sundaresan, Zamboni, Dinsdale, et al., 2021; Sundaresan et al., 2022), a deep-learning tool that utilises the spatial distribution of WMH in the loss functions during optimisation. Resulting maps were thresholded at 0.05, binarised and masked with subject-specific WM masks to remove false positives outside of the white matter. Maps then underwent visual inspection and comparison with overlayed FLAIR images. On the basis of the resulting segmented WMH maps, the confluence metric Conf(*A*) was calculated for each subject, providing a score between 0 and 1. WMH volumes were determined as the sum of all voxels in the WMH map multiplied with voxel size in mm. A log base 10 transformation of WMH volumes was performed as recommended previously (Roseborough et al., 2023) in order to account for the right-skewed distribution of lesion volume.

### 3.6 Separate confluence scores for periventricular and deep WMH

Based on previous studies stating that periventricular and deep WMH (PWMH/DWMH) might be different pathological entities (Fazekas et al., 1998; R. Schmidt, Schmidt, et al., 2011; Armstrong et al., 2020; Voorter et al., 2025) and should be reported separately (Roseborough et al., 2023), we calculated distinct PWMH and DWMH confluence scores for each participant. An additional reason for the distinct scores was that we aimed to compare confluence scores with Fazekas scores which are determined for deep and periventricular WM separately. There is no consensus as to how the two areas should be separated and different methods have been suggested. We chose the method of a 10 mm distance from the ventricle edges since it is commonly used and was recommended previously (Griffanti et al., 2018). To this end, ventricle distance maps created within the TrUE-Net preprocessing pipeline were thresholded so that periventricular WM was defined as WM within a 10 mm distance from the ventricles, and deep WM was defined as WM further than 10 mm away.

### 3.7 WMH rating with Fazekas scale

WMH burden was additionally assessed by ratings using the Fazekas scale (Fazekas et al., 1987). This visual scale is a widely used WMH rating system and it separately takes into account periventricular and deep white matter, assigning grades between 0 and 3 depending on the extent of lesions. The rater was blind to confluence scores and participant diagnosis. Figure 2 shows images representative of the different grades.

**Figure 2:**
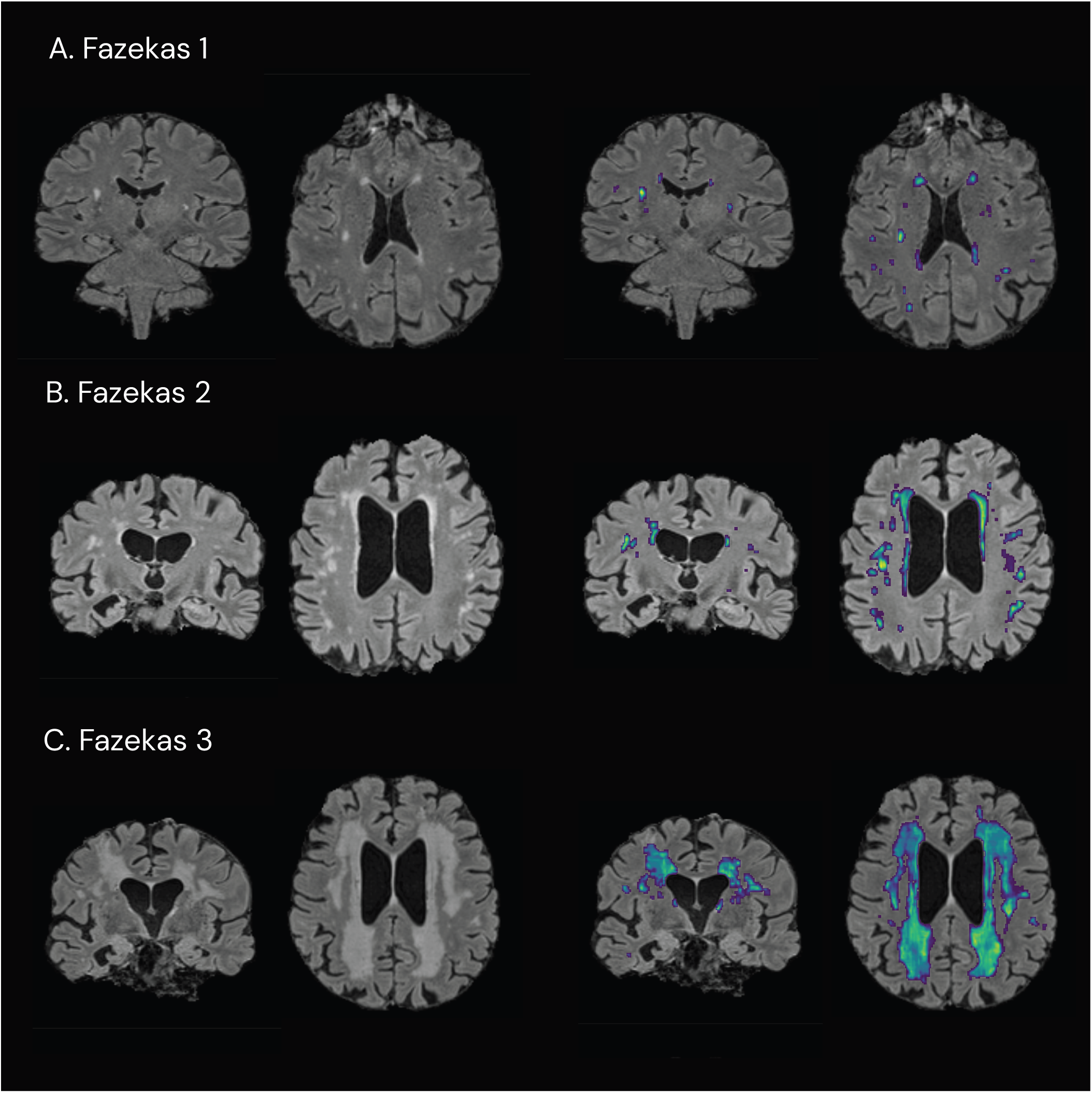
Examples of FLAIR MRI images with different Fazekas scores. The images on the right show WMH maps generated by TrUE-Net segmentation. The WMH become more confluent with increasing Fazekas score.

### 3.8 Statistical analyses

All analyses were conducted in Python (version 3.14.1). False discovery rate (FDR; Benjamini and Hochberg, 1995) adjustment was applied where appropriate to correct resulting *p*-values for multiple comparisons.

Pearson’s correlation coefficient was used to assess the linear association between WMH volume and confluence. Next, the association of the confluence metric in deep, periventricular and whole-brain WM with the corresponding Fazekas scores was tested with Pearson’s correlation coefficient. In order to explore how WMH properties change with age, linear regressions of confluence and volume respectively against age were conducted.

To examine the relationship between WMH confluence and cognitive performance, we then ran separate linear regressions of the score of each ACE subtest (memory, attention, verbal fluency, visuospatial abilities, language) as well as the total ACE score against WMH confluence, estimated total intracranial volume, age and sex (Model 1). In an additional regression model (Model 2), WMH volume and the interaction of volume and confluence were included as predictors in order to test whether WMH confluence had an effect on test scores above and beyond the effect of volume. In both regression models, the predictors volume and confluence were mean-centred to improve interpretability, but predictors were not standardised in order to preserve the natural units. Some of the ACE subscores displayed a ceiling effect (full marks in the attention, language and visuospatial abilities subtests were achieved by 28%, 27% and 35% of subjects who completed the ACE, respectively). The ceiling effect leads to violation of one of the assumptions of ordinary least squares regression, namely linearity of conditional expectation. Therefore we tested the relationship between attention, language and visuospatial abilities scores and confluence with a Tobit regression with the R package censReg (Henningsen, 2010), including the same covariates as in the linear regressions of the other ACE subtests.

Finally, in order to test whether WMH confluence differed between diagnostic groups, Welch’s *t*-test was performed for all possible pairs of diagnoses.

All statistical analyses were performed for the whole-brain WMH confluence scores as well as for DWMH and PWMH confluence scores separately.

### 3.9 Validation dataset

In order to validate our findings, we used an independent dataset which had not been involved in developing the metric, the DZNE-Longitudinal Cognitive Impairment and Dementia Study (DELCODE; Jessen et al., 2018). DELCODE is an observational longitudinal memory clinic study based in Germany for the purpose of investigating the early preclinical stage of AD, particularly in patients with subjective cognitive decline. Accordingly, the diagnoses represented in DELCODE are Alzheimer’s disease, mild cognitive impairment and subjective cognitive decline. Additionally, cognitively normal participants are part of the study. In total, 893 individuals were included in the analysis.

Cognitive performance was assessed on five cognitive domain scores (learning and memory, language ability, executive functions and mental processing speed, working memory, and visuospatial abilities). These are compound scores that were derived from a confirmatory factor analysis of a larger test battery (Wolfsgruber et al., 2020). The overall score of that test battery was also reported (NPT global score). Additionally, the Preclinical Alzheimer Cognitive Composite (PACC; Donohue et al., 2014) was administered.

WMH were segmented with the AI-augmented version of the Lesion Segmentation Toolbox (LST-AI; Wiltgen et al., 2024) using T1-weighted and FLAIR images. Fazekas rating was performed by an expert rater, WMH volume and confluence were determined as in the QMIN-MC cohort.

Repeating the statistical analysis in the QMIN-MC cohort, we conducted a Pearson correlation analysis between confluence and WMH volume as well as between confluence and Fazekas scores, a linear regression of each cognitive score against WMH confluence and covariates of no interest, and a *t*-test of confluence scores for each pair of diagnostic groups.

## 4 Results

### 4.1 Overview

We analysed data from the Quantitative MRI in NHS Memory Clinics dataset (QMIN-MC; *n* = 1073, mean age = 72); demographic details are shown in Table 1. Our novel quantitative metric of WMH confluence was applied to the imaging data and we explored whether confluence was associated with age, cognitive performance and Fazekas scores, and whether it differed between diagnostic groups. Whole-brain WMH confluence scores across all diagnostic groups ranged from 0.02 to 0.53 (1 being the score of a hypothetical image consisting entirely of WMH voxels), with a mean (SD) of 0.20 (0.10). Deep WMH confluence scores ranged between 0.00 and 0.50 (mean 0.11, SD 0.06), periventricular WMH confluence scores ranged between 0.02 and 0.57 (mean 0.21, SD 0.12).

**Table 1:** Demographics and sample characteristics by diagnostic group. QMIN-MC sample*^a^*.

|  | Overall | AD | Non-AD dem. | Vascular dem. | Mixed dem. | MCI | Other | No CI | p-Value |
| --- | --- | --- | --- | --- | --- | --- | --- | --- | --- |
| n | 1073 | 283 | 28 | 29 | 53 | 141 | 362 | 177 |  |
| Age, mean (SD) | 73 (11) | 75 (9) | 72 (9) | 78 (9) | 79 (7) | 76 (7) | 74 (9) | 59 (11) | **<0.001\*\*\*** |
| Gender, n (%) |  |  |  |  |  |  |  |  | 0.295 |
| Female | 529 (49.3) | 163 (57.6) | 12 (42.9) | 13 (44.8) | 25 (47.2) | 66 (46.8) | 169 (46.7) | 81 (45.8) |  |
| Male | 537 (50.0) | 118 (41.7) | 16 (57.1) | 16 (55.2) | 28 (52.8) | 74 (52.5) | 189 (52.2) | 96 (54.2) |  |
| Other | 7 (0.7) | 2 (0.7) | 0 (0.0) | 0 (0.0) | 0 (0.0) | 1 (0.7) | 4 (1.1) | 0 (0.0) |  |
| Years of education, mean (SD) | 12.3 (3.2) | 12.0 (2.9) | 12.5 (2.7) | 10.8 (1.7) | 12.3 (3.2) | 12.6 (3.4) | 12.2 (3.1) | 13.0 (3.5) | **0.005\*\*** |
| Whole-brain WMH volume, mean (SD) | 11239 (14637) | 12086 (14137) | 11938 (15029) | 32040 (26814) | 18935 (16745) | 13710 (16010) | 10998 (13540) | 2989 (4038) | **<0.001\*\*\*** |
| Whole-brain WMH confluence, mean (SD) | 0.200 (0.100) | 0.216 (0.087) | 0.215 (0.098) | 0.311 (0.124) | 0.262 (0.103) | 0.218 (0.105) | 0.201 (0.098) | 0.124 (0.058) | **<0.001\*\*\*** |
| DWMH volume, mean (SD) | 2672 (5557) | 2730 (5673) | 2122 (4249) | 9862 (11906) | 3928 (4993) | 3505 (6314) | 2576 (5278) | 730 (1513) | **<0.001\*\*\*** |
| DWMH confluence, mean (SD) | 0.114 (0.062) | 0.114 (0.060) | 0.113 (0.063) | 0.181 (0.093) | 0.138 (0.060) | 0.125 (0.067) | 0.114 (0.061) | 0.086 (0.039) | **<0.001\*\*\*** |
| PWMH volume, mean (SD) | 8222 (9934) | 8939 (9345) | 9413 (10970) | 21668 (16030) | 14550 (12902) | 9814 (10538) | 8079 (9313) | 2123 (2892) | **<0.001\*\*\*** |
| PWMH confluence, mean (SD) | 0.212 (0.115) | 0.229 (0.101) | 0.225 (0.114) | 0.346 (0.136) | 0.277 (0.116) | 0.233 (0.120) | 0.214 (0.114) | 0.123 (0.066) | **<0.001\*\*\*** |
| Total ACE (0-100), mean (SD) | 73.4 (16.4) | 63.7 (15.3) | 75.2 (16.4) | 62.3 (19.2) | 66.8 (13.4) | 77.6 (10.7) | 75.1 (16.4) | 84.9 (12.2) | **<0.001\*\*\*** |
| Attention (0-18), mean (SD) | 14.8 (3.5) | 12.9 (3.9) | 16.2 (3.5) | 12.7 (3.9) | 13.4 (3.8) | 15.9 (2.1) | 15.1 (3.3) | 17.0 (1.9) | **<0.001\*\*\*** |
| Memory (0-26), mean (SD) | 15.3 (6.1) | 11.3 (4.9) | 17.0 (6.1) | 14.4 (6.2) | 11.8 (5.0) | 16.5 (5.2) | 16.3 (6.1) | 19.2 (5.2) | **<0.001\*\*\*** |
| Fluency, mean (SD) | 33.7 (8.8) | 29.4 (7.5) | 32.3 (8.9) | 27.4 (10.7) | 30.6 (7.3) | 34.3 (6.5) | 34.1 (9.0) | 40.6 (7.5) | **<0.001\*\*\*** |
| Language (0-26), mean (SD) | 22.9 (4.0) | 21.6 (4.6) | 22.2 (4.2) | 19.6 (7.1) | 23.2 (3.0) | 23.9 (2.9) | 23.1 (4.0) | 24.1 (2.8) | **<0.001\*\*\*** |
| Visuospatial ab. (0-16), mean (SD) | 13.7 (2.8) | 12.6 (3.5) | 13.5 (3.3) | 11.6 (4.0) | 13.4 (2.6) | 14.6 (1.4) | 13.9 (2.6) | 15.0 (1.8) | **<0.001\*\*\*** |

<sup>a</sup> AD = Alzheimer's dementia; Non-AD dem. = non-Alzheimer's disease dementia; No CI = No cognitive impairment; WMH = white matter hyperintensity; DWMH = deep white matter hyperintensity; PWMH = periventricular white matter hyperintensity; ACE = Addenbrooke's Cognitive Examination. *P*-values have been FDR-corrected for multiple comparisons, bold *p*-values are significant. Asterisks indicate significance levels: *p*\* < 0.05, *p*\*\* < 0.01, *p*\*\*\* < 0.001.

### 4.2 Confluence scores correlate strongly with log-transformed WMH volume and with Fazekas scores

Confluence scores correlated strongly with WMH volume after volume log-transformation (*r* = 0.91 for DWMH; *r* = 0.80 for PWMH; *r* = 0.92 for whole-brain WMH; Figure 3). Furthermore, confluence scores correlated strongly with Fazekas scores (*r* = 0.65 for DWMH; *r* = 0.80 for PWMH; *r* = 0.80 for whole-brain WMH; Figure 4). Correlations between raw WMH volume and Fazekas scores were lower (*r* = 0.56 for DWMH; *r* = 0.65 for PWMH; *r* = 0.70 for whole-brain WMH; data not shown), but log-transformed WMH volume correlated similarly with Fazekas scores (*r* = 0.80 for DWMH; *r* = 0.83 for PWMH; *r* = 0.87 for whole-brain WMH; data not shown).

**Figure 3:**
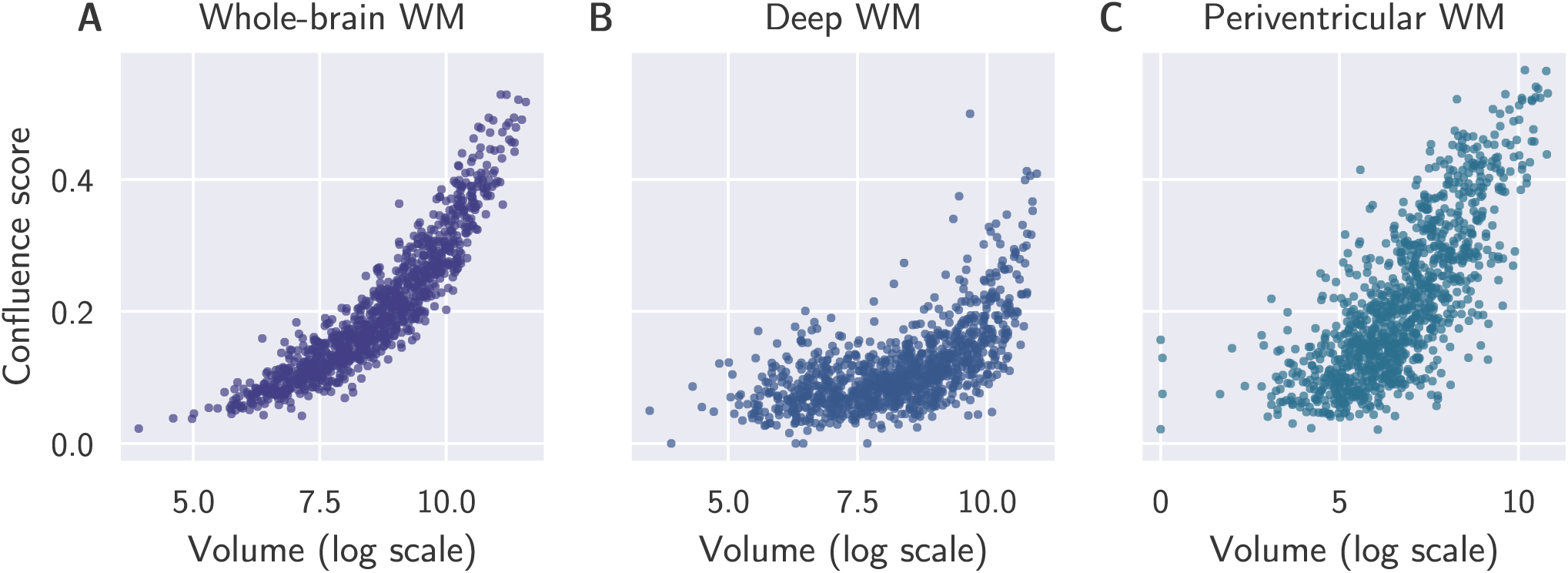
Relationship between WMH confluence and volume in whole-brain WM (A), deep WM (B) and periventricular WM (C). QMIN-MC, *N* = 1073.

**Figure 4:**
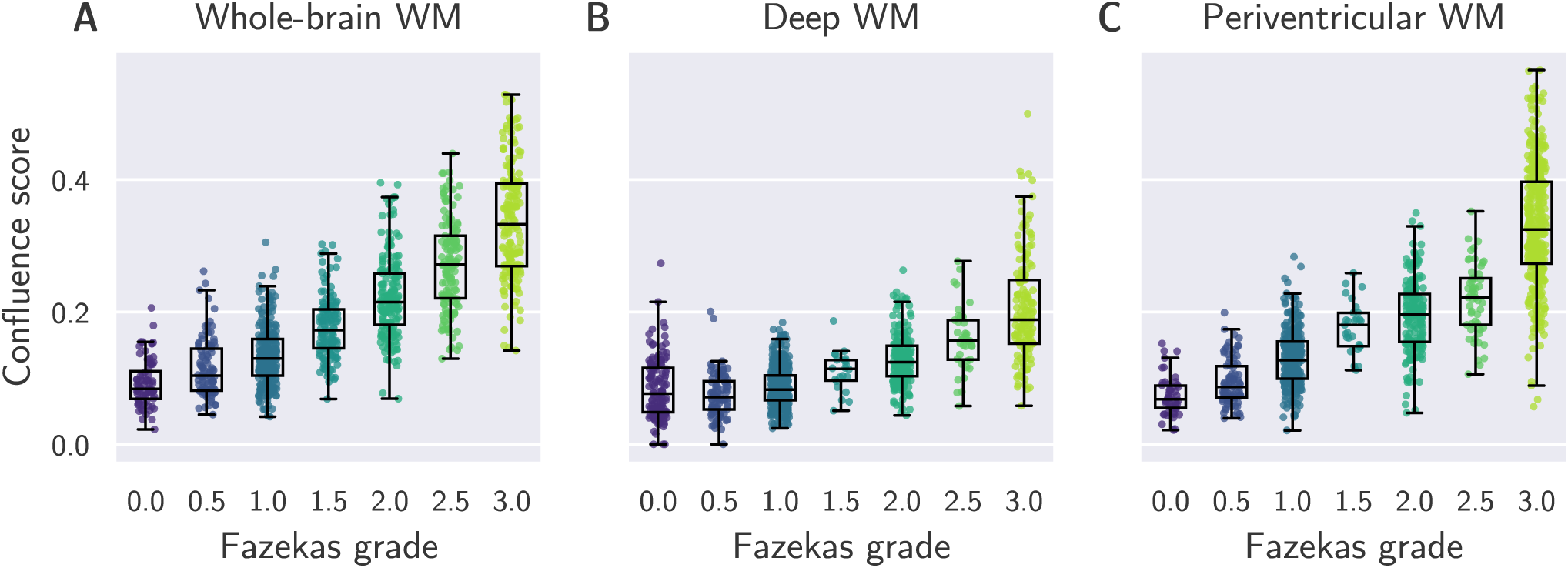
Relationship between Fazekas scores and WMH confluence in whole-brain WM (A), deep WM (B) and periventricular WM (C). QMIN-MC, *N* = 1073.

### 4.3 Confluence of whole-brain, deep and periventricular WMH increases with age

Age was associated with higher confluence scores (*β* = .005, *t* = 18.95, *p <* 0.001, *R*^2^ = 0.26) and greater WMH volume (*β* = 526.08, *t* = 13.734, *p <* 0.001, *R*^2^ = 0.15) across the whole brain. An age difference of 10 years corresponded to a WMH confluence difference of 0.05, which is a difference of less than one standard deviation (Figure 5). Similarly, an age association was found for confluence of deep and periventricular WMH (DWMH: *β* = −0.002, *t* = 9.26, *p <* 0.001, *R*^2^ = 0.08; PWMH: *β* = 0.005, *t* = 19.37, *p <* 0.001, *R*^2^ = 0.27) as well as for their volume (DWMH: *β* = 119.58, *t* = 15.37, *p <* 0.001, *R*^2^ = 0.06; PWMH: *β* = 394.53, *t* = 15.49, *p <* 0.001, *R*^2^ = 0.19; Figure 5). An age difference of 10 years corresponded to a difference in DWMH confluence of 0.02 and a difference in PWMH confluence of 0.05, both differences are less than one SD.

**Figure 5:**
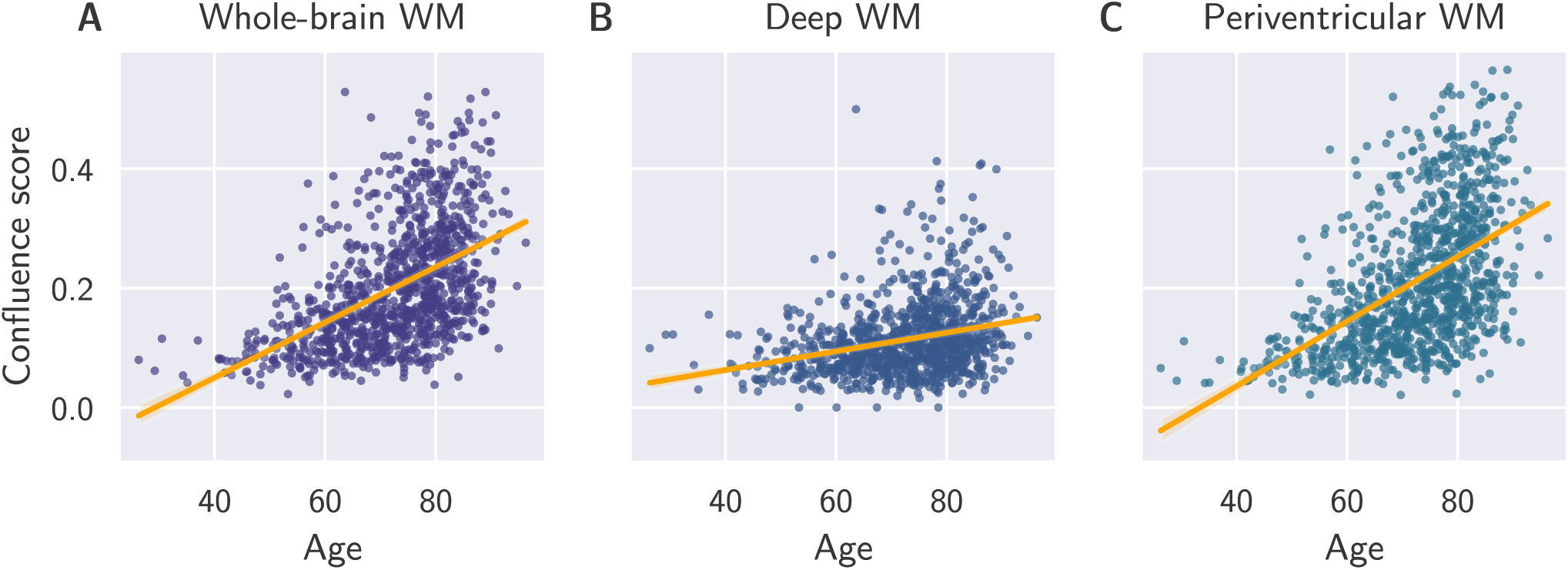
WMH confluence was associated with age, for lesions in whole-brain WM (A), deep WM (B) and periventricular WM (C). QMIN-MC, *N* = 1073.

### 4.4 Confluence is associated with performance across cognitive domains, but does not explain variance in addition to WMH volume

Regressions using Model 1, which included WMH confluence and covariates of no interest as predictors, showed a significant effect of WMH confluence on all ACE subscales (attention, memory, fluency, language and visuospatial abilities) as well as the total ACE score (Table 2 and Figure 6; s. Table **??** in Supplementary Materials for the full regression results). A difference in the whole-brain confluence score of 0.1 corresponded to a decrease in the total ACE score of 3.5 out of 100 points, a decrease in the attention score of 0.6 out of 18, a decrease in the memory score of 1.2 out of 26, a decrease in fluency of 1.6 words, a decrease in the language score of 1.0 out of 26 points and a decrease in the visuospatial function score of 0.4 out of 26 points. Periventricular WMH confluence was also linked to performance in all five ACE subtests and the total score, while deep WMH confluence was associated with performance in the memory, fluency and language subtests as well as the total ACE score. Similar results were obtained when additionally including raw WMH volume in the model (results not shown), with no significant effect of raw volume on cognitive performance. However, after additionally adjusting for *log-transformed* WMH volume in Model 2, WMH confluence did not explain variance in cognitive performance beyond that accounted for by WMH volume. Instead, only log-transformed periventricular WMH volume had an effect on fluency and language (Table 3; s. Table **??** in Supplementary Materials for full regression results).

**Figure 6:**
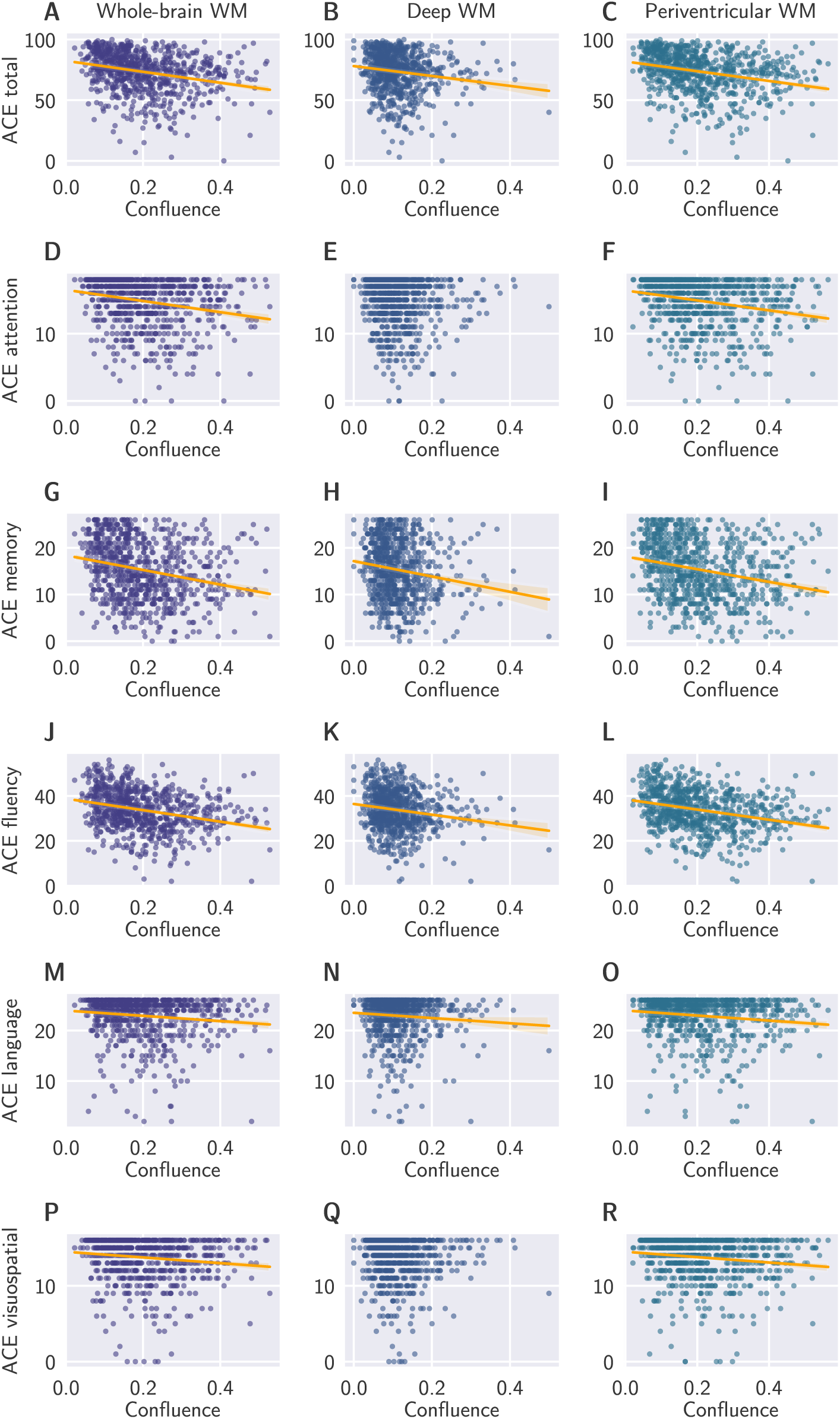
Poor performance in cognitive domains of the ACE was associated with both deep and periventricular white matter confluence. Regression lines are shown for relationships that were significant using Model 1. QMIN-MC, *N* = 1073.

**Table 2:** Relationship between ACE subtests and WMH: Regression Model 1 with confluence as a predictor. QMIN-MC data*^a^*.

| Outcome variable | Whole-brain WMH confluence |  |  |  |  | DWMH confluence |  |  |  |  | PWMH confluence |  |  |  |  |
| --- | --- | --- | --- | --- | --- | --- | --- | --- | --- | --- | --- | --- | --- | --- | --- |
| | $\beta$ | SD | t | $p_{FDR}$ | $R^2$ | $\beta$ | SD | t | $p_{FDR}$ | $R^2$ | $\beta$ | SD | t | $p_{FDR}$ | $R^2$ |
| Total ACE | -35.07 | 6.71 | -5.23 | < **0.001** *** | 0.09 | -25.44 | 9.71 | -2.62 | **0.012** * | 0.07 | -31.68 | 5.83 | -5.43 | < **0.001** *** | 0.10 |
| Attention | -6.86 | 1.86 | -3.69 | < **0.001** *** |  | -1.97 | 2.67 | -0.74 | 0.487 |  | -6.11 | 1.62 | -3.77 | < **0.001** *** |  |
| Memory | -11.78 | 2.53 | -4.66 | < **0.001** *** | 0.08 | -11.00 | 3.63 | -3.03 | **0.004** ** | 0.06 | -10.21 | 2.20 | -4.64 | < **0.001** *** | 0.08 |
| Fluency | -16.33 | 3.60 | -4.54 | < **0.001** *** | 0.11 | -12.64 | 5.16 | -2.45 | **0.018** * | 0.09 | -14.67 | 3.13 | -4.69 | < **0.001** *** | 0.11 |
| Language | -9.66 | 2.13 | -4.54 | < **0.001** *** |  | -8.29 | 3.03 | -2.74 | **0.009** ** |  | -8.88 | 1.85 | -4.80 | < **0.001** *** |  |
| Visuospatial ab. | -3.77 | 1.71 | -2.21 | **0.031** * |  | 0.51 | 2.45 | 0.21 | 0.835 |  | -3.86 | 1.48 | -2.61 | **0.012** * |  |

<sup>a</sup> Predictors included in the model: WMH confluence, age, sex, total intracranial volume. $P$ -values have been FDR-corrected for 18 tests, bold $p$ -values are significant. Asterisks indicate significance levels: $p^* < 0.05$ , $p^{**} < 0.01$ , $p^{***} < 0.001$ . No $R^2$ is available for Tobit regressions (attention, language and visuospatial abilities).

**Table 3:**
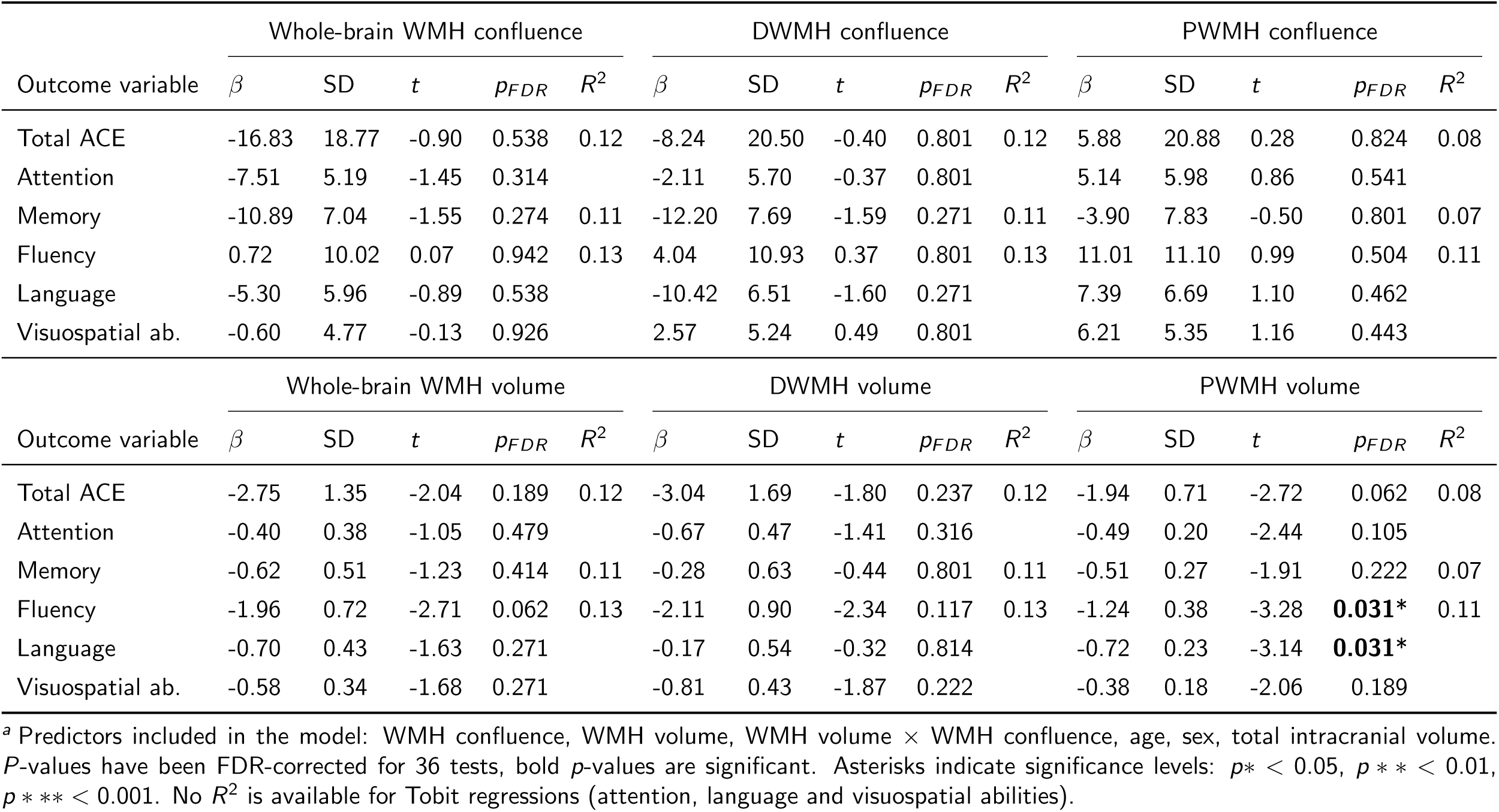
Relationship between ACE subtests and WMH: Regression Model 2 with confluence and volume as predictors. QMIN-MC data*^a^*.

### 4.5 Confluence of whole-brain, deep and periventricular WMH differs between diagnostic groups

Welch’s *t*-test for pairs of diagnostic groups identified significant differences in WMH confluence between participants with vascular and mixed dementia and all other diagnostic groups (Table 4; Figure 7). Moreover, participants with no cognitive impairment displayed lower whole-brain WMH confluence than any other diagnostic group. With few exceptions these findings were the same for deep and periventricular WMH. Confluence scores of the remaining diagnostic groups were largely overlapping and could not be distinguished.

**Figure 7:**
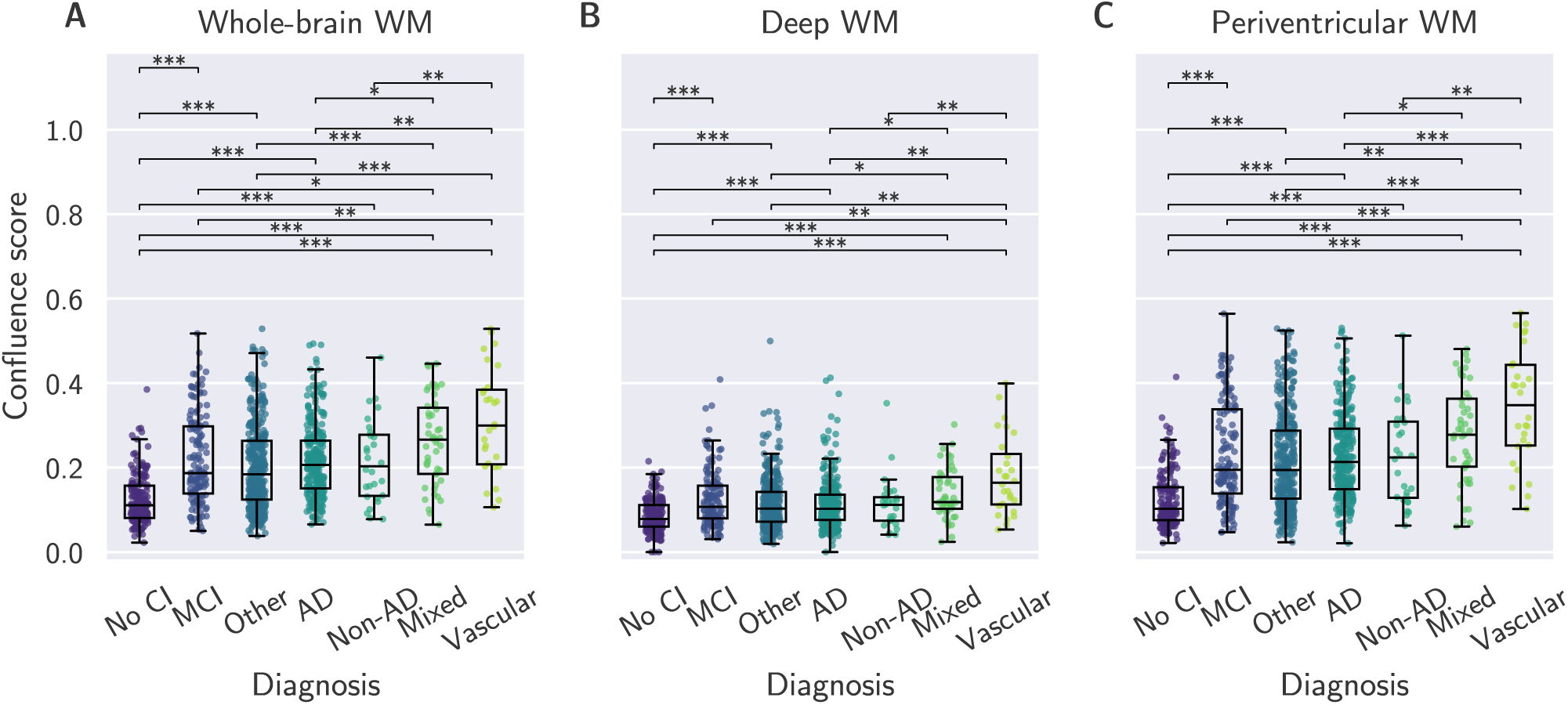
WMH confluence in whole-brain (A), deep (B) and periventricular WM (C), by diagnostic group. Participants with vascular dementia and with no cognitive impairment differed in WMH confluence from almost all other diagnostic groups. QMIN-MC, *N* = 1073. Control = no cognitive impairment; MCI = mild cognitive impairment; AD = Alzheimer’s disease; Non-AD = other non-AD dementia; Mixed = mixed dementia; Vascular = vascular dementia.

**Table 4:** Results of Welch’s *t*-test comparing WMH confluence between pairs of diagnostic groups. QMIN-MC data*^a^*.

| Diagnosis pair | Whole-brain WMH confluence |  | DWMH confluence |  | PWMH confluence |  |
| --- | --- | --- | --- | --- | --- | --- |
| | $t$ | $pFDR$ | $t$ | $pFDR$ | $t$ | $pFDR$ |
| AD vs. Non-AD dementia | 0.19 | 0.882 | 0.25 | 0.868 | 0.28 | 0.862 |
| AD vs. Mixed dementia | -3.18 | **0.005\*\*** | -2.96 | **0.008\*\*** | -2.96 | **0.008\*\*** |
| AD vs. Vascular dementia | -3.44 | **0.004\*\*** | -3.39 | **0.005\*\*** | -3.85 | **0.002\*\*** |
| AD vs. Other | 2.03 | 0.065 | -0.10 | 0.927 | 1.76 | 0.115 |
| AD vs. MCI | -0.09 | 0.927 | -1.55 | 0.160 | -0.22 | 0.868 |
| AD vs. No cognitive impairment | 13.85 | <**0.001\*\*\*** | 6.52 | <**0.001\*\*\*** | 13.77 | <**0.001\*\*\*** |
| Non-AD dementia vs. Mixed dementia | -2.26 | **0.046\*** | -2.16 | 0.054 | -2.16 | 0.054 |
| Non-AD dementia vs. Vascular dementia | -2.90 | **0.010\*\*** | -3.01 | **0.008\*\*** | -3.25 | **0.005\*\*** |
| Non-AD dementia vs. Other | 0.60 | 0.655 | -0.30 | 0.862 | 0.40 | 0.808 |
| Non-AD dementia vs. MCI | -0.22 | 0.868 | -1.04 | 0.371 | -0.38 | 0.812 |
| Non-AD dementia vs. No cognitive impairment | 4.82 | <**0.001\*\*\*** | 2.24 | 0.053 | 4.61 | <**0.001\*\*\*** |
| Mixed dementia vs. Vascular dementia | -1.23 | 0.279 | -1.58 | 0.160 | -1.66 | 0.143 |
| Mixed dementia vs. Other | 4.15 | <**0.001\*\*\*** | 2.94 | **0.009\*\*** | 3.82 | <**0.001\*\*\*** |
| Mixed dementia vs. MCI | 2.84 | **0.010\*\*** | 1.75 | 0.119 | 2.56 | **0.020\*** |
| Mixed dementia vs. No cognitive impairment | 9.43 | <**0.001\*\*\*** | 6.23 | <**0.001\*\*\*** | 9.25 | <**0.001\*\*\*** |
| Vascular dementia vs. Other | 4.06 | <**0.001\*\*\*** | 3.37 | **0.005\*\*** | 4.41 | <**0.001\*\*\*** |
| Vascular dementia vs. MCI | 3.26 | **0.005\*\*** | 2.73 | **0.018\*** | 3.60 | **0.003\*\*** |
| Vascular dementia vs. No cognitive impairment | 7.38 | <**0.001\*\*\*** | 5.15 | <**0.001\*\*\*** | 7.90 | <**0.001\*\*\*** |
| Other vs. MCI | -1.57 | 0.160 | -1.51 | 0.167 | -1.52 | 0.167 |
| Other vs. No cognitive impairment | 11.71 | <**0.001\*\*\*** | 6.92 | <**0.001\*\*\*** | 11.87 | <**0.001\*\*\*** |
| MCI vs. No cognitive impairment | 9.77 | <**0.001\*\*\*** | 6.48 | <**0.001\*\*\*** | 9.89 | <**0.001\*\*\*** |

<sup>a</sup> AD = Alzheimer's dementia; MCI = mild cognitive impairment; Non-AD dementia = non-Alzheimer's disease dementia; Control = participants with functional/attentional memory symptoms; DWMH = deep white matter hyperintensity; PWMH = periventricular white matter hyperintensity. $P$ -values have been FDR-corrected for 21 tests, bold $p$ -values are significant. Asterisks indicate significance levels: $p^* < 0.05$ , $p^{**} < 0.01$ , $p^{***} < 0.001$ .

### 4.6 3D confluence analysis is equivalent and more computationally intense than 2D analysis

We repeated the analysis with the 3D confluence score metrics. There was a high correlation between 3D and averaged 2D confluence metrics of *r* = 0.96, indicating that differences between the two measures were negligible (Figure 8). Results of statistical analyses did not change when using 3D confluence scores as opposed to 2D. Furthermore, the 2D option has lower computational requirements: Computational complexity of the 2D slice formula is O(*d* ^4^), averaging over 2D slices then has computational complexity O(*d* ^5^). The 3D formula on the other hand has computational complexity O(*d* ^6^). Therefore we focused on implementing the computationally more tractable 2D option.

**Figure 8:**
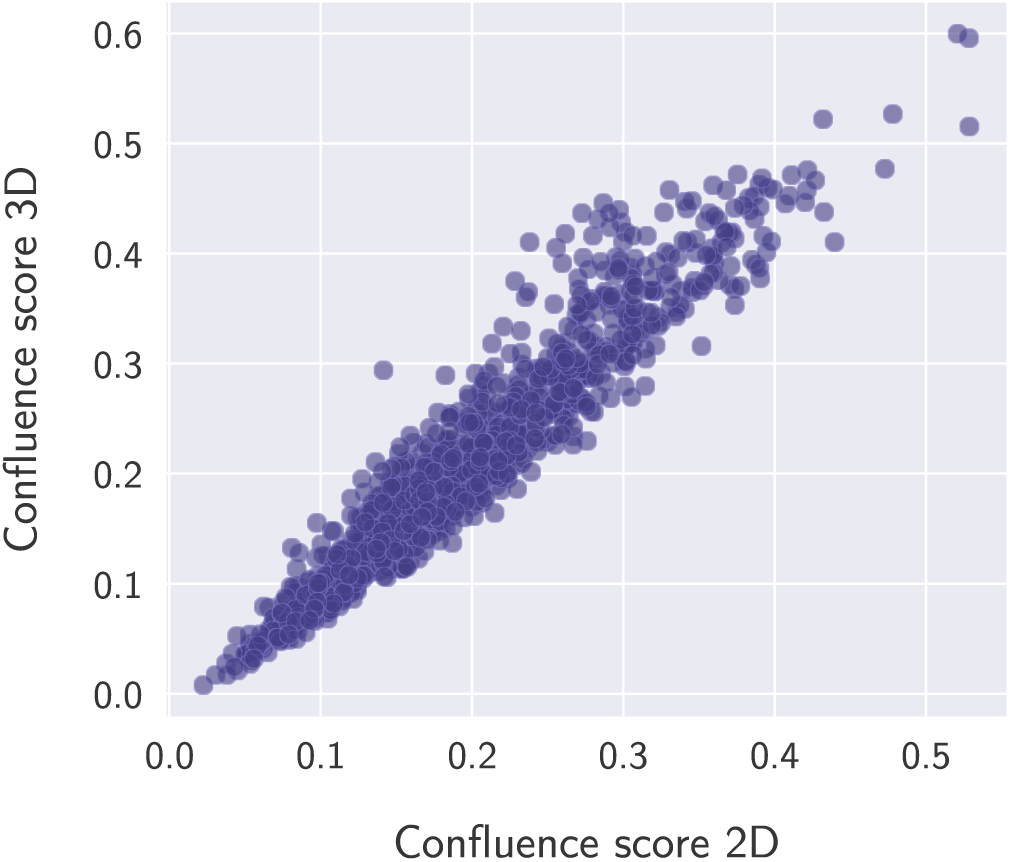
WMH confluence metrics calculated in 2D and in 3D show a strong correlation.

### 4.7 Validation in an independent dataset

We used the DELCODE cohort for validation (*n* = 893, mean age = 71; demographic details are shown in Table 5. In the DELCODE group, a similar but slightly smaller more narrow range of confluence scores was found as compared to the QMIN-MC group. The correlations between confluence scores and Fazekas scores were moderate (*r* = 0.63 for whole-brain WMH; *r* = 0.31 for DWMH; *r* = 0.56 for PWMH; Fig. 9). For whole-brain and periventricular WMH confluence, we found an association with all cognitive domains including memory, executive function, visuospatial abilities, language, working memory, the NPT global score and the PACC score (Figure 11). For deep WMH, we identified an effect of confluence on memory and language.

**Figure 9:**
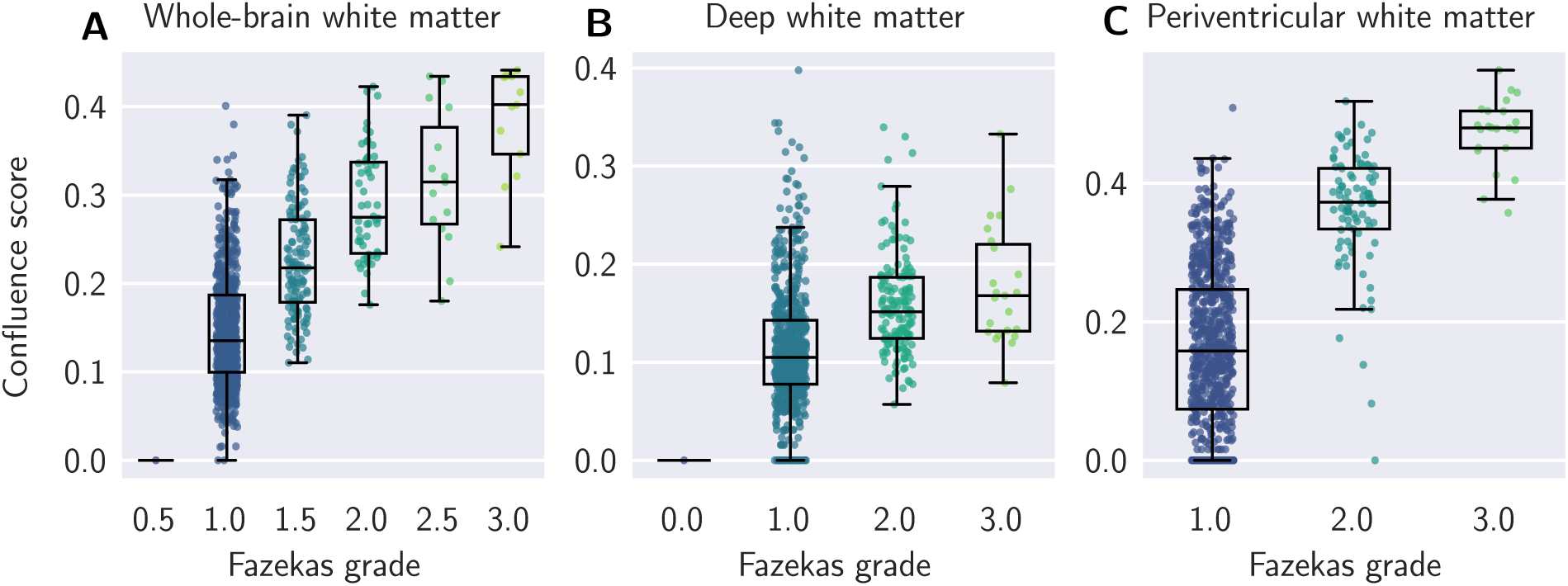
WMH confluence scores are correlated with Fazekas scores. Left: Whole-brain WMH; middle: Deep WMH; right: Periventricular WMH. DELCODE, *N* = 893.

**Figure 10:**
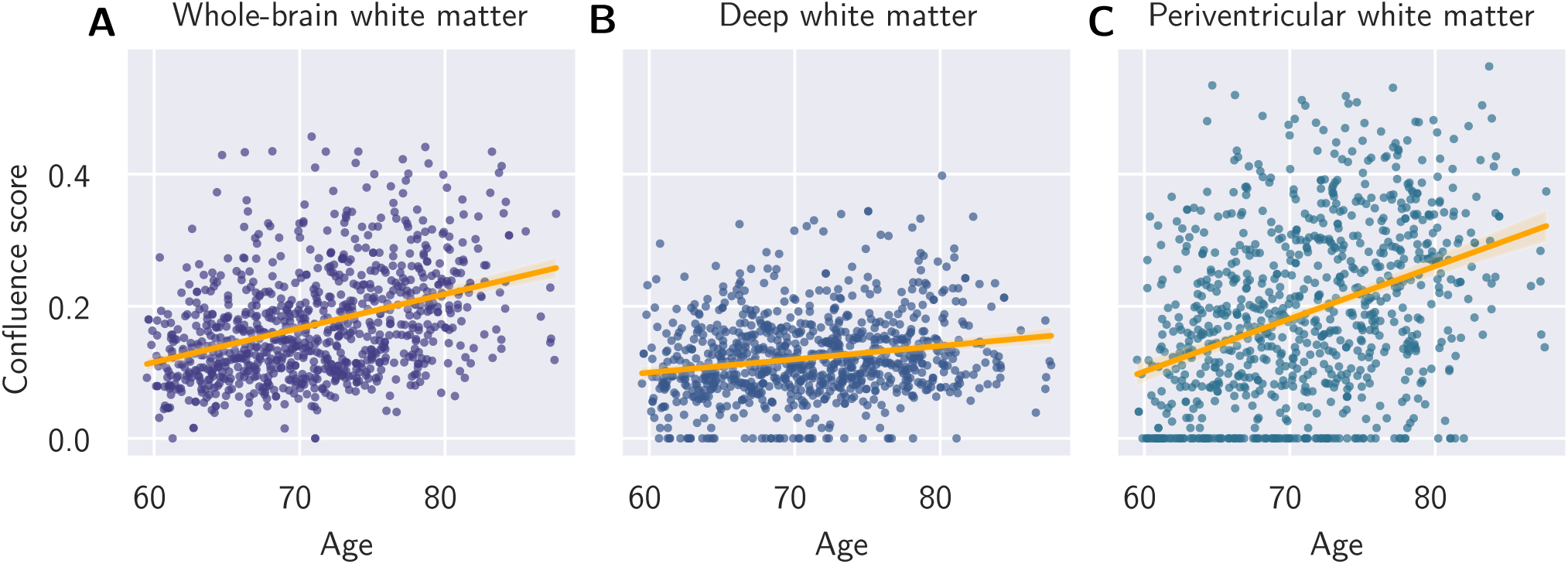
Confluence of whole-brain, deep and periventricular WMH increases with age. DELCODE, *N* = 893.

**Figure 11:**
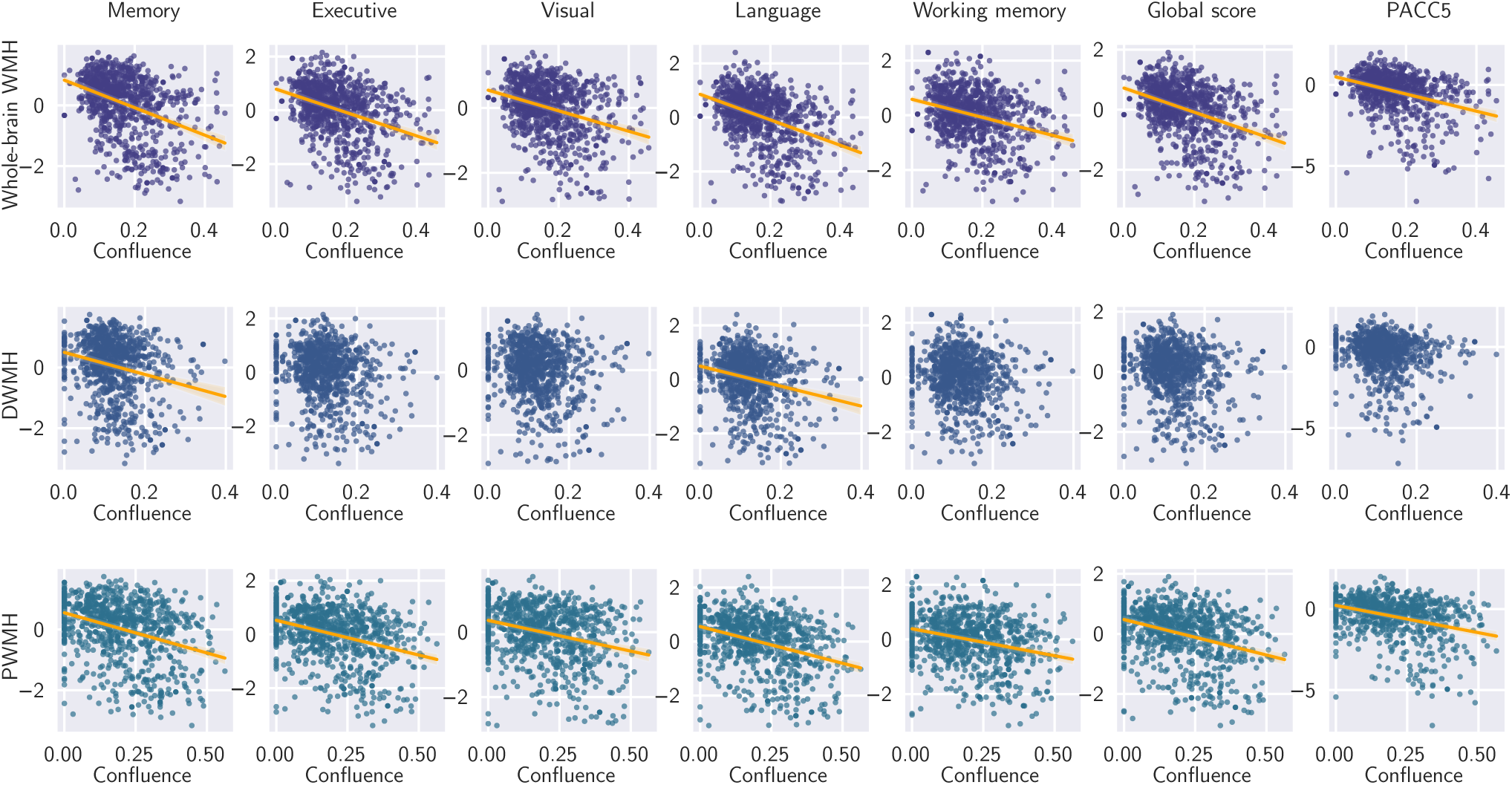
Relationship between cognitive factor scores and WMH confluence. Whole-brain and periventricular WMH confluence were associated with cognitive performance across domains while deep WMH confluence showed no association with performance in any tested cognitive domain. DELCODE, *N* = 893.

**Figure 12:**
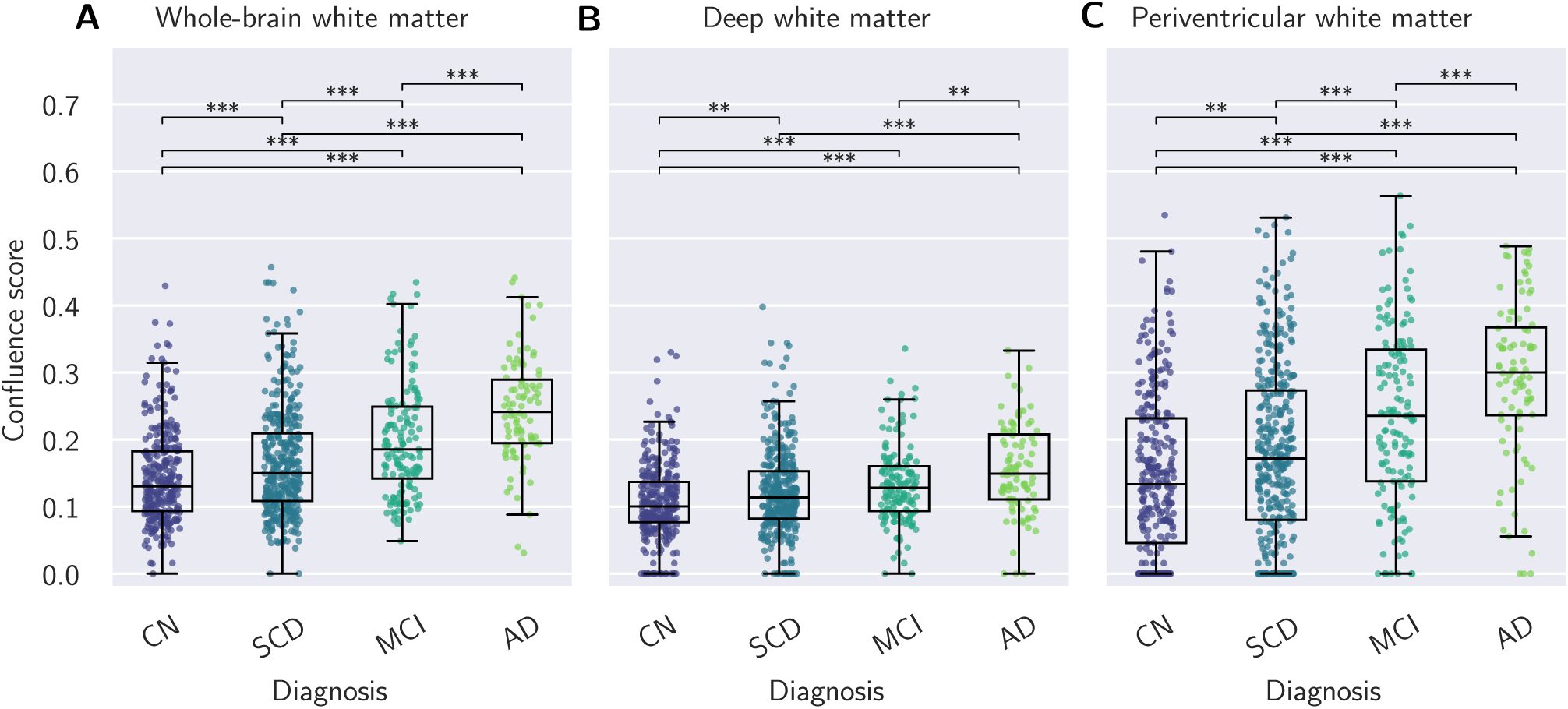
WMH confluence differed between most diagnostic groups in the DELCODE cohort. DELCODE, *N* = 893. CN = control; SCD = subjective cognitive decline; MCI = mild cognitive impairment; AD = Alzheimer’s disease.

**Table 5:** Demographics and sample characteristics by diagnostic group. DELCODE data*^a^*.

|  | No CI | SCD | MCI | AD | p-value |
| --- | --- | --- | --- | --- | --- |
| n | 294 | 364 | 143 | 92 |  |
| Age | 68.70 (5.36) | 71.18 (6.17) | 73.23 (5.71) | 74.97 (6.31) | < **0.001** *** |
| Gender, n (%) |  |  |  |  |  |
| Female | 175 (59.5) | 168 (46.2) | 69 (48.3) | 51 (55.4) |  |
| Male | 119 (40.5) | 196 (53.8) | 74 (51.7) | 41 (44.6) |  |
| Years of education | 14.63 (2.75) | 14.99 (3.03) | 13.90 (3.07) | 13.01 (3.29) | < **0.001** *** |
| Whole-brain WMH volume | 2914.84 (5561.25) | 4577.60 (7873.36) | 7416.29 (11146.55) | 10612.45 (9673.18) | < **0.001** *** |
| Whole-brain WMH confluence | 0.14 (0.07) | 0.17 (0.08) | 0.20 (0.08) | 0.24 (0.08) | < **0.001** *** |
| PWMH volume | 1779.95 (4712.53) | 3061.77 (6834.30) | 5487.20 (10145.86) | 7450.33 (8481.52) | < **0.001** *** |
| PWMH confluence | 0.15 (0.12) | 0.18 (0.13) | 0.24 (0.13) | 0.29 (0.12) | < **0.001** *** |
| DWMH volume | 763.06 (1224.03) | 1060.84 (1630.81) | 1373.69 (1861.69) | 2331.92 (2718.00) | < **0.001** *** |
| DWMH confluence | 0.11 (0.06) | 0.12 (0.06) | 0.13 (0.06) | 0.16 (0.07) | < **0.001** *** |
| Memory | 0.60 (0.46) | 0.41 (0.56) | -0.73 (0.69) | -1.95 (0.52) | < **0.001** *** |
| Executive function | 0.51 (0.55) | 0.33 (0.69) | -0.49 (0.77) | -1.83 (0.78) | < **0.001** *** |
| Language | 0.54 (0.49) | 0.38 (0.59) | -0.62 (0.64) | -1.88 (0.66) | < **0.001** *** |
| Visuospatial ab. | 0.33 (0.52) | 0.27 (0.61) | -0.44 (0.66) | -1.47 (0.78) | < **0.001** *** |
| Working memory | 0.35 (0.56) | 0.31 (0.70) | -0.41 (0.76) | -1.53 (0.78) | < **0.001** *** |
| Global score | 0.47 (0.44) | 0.34 (0.55) | -0.54 (0.61) | -1.73 (0.62) | < **0.001** *** |
| PACC5 | 0.17 (0.58) | -0.08 (0.68) | -1.38 (0.98) | -3.75 (1.26) | < **0.001** *** |

<sup>a</sup> No CI = no cognitive impairment; SCD = subjective cognitive decline; MCI = mild cognitive impairment; AD = Alzheimer's disease. *P*-values have been FDR-corrected for multiple comparisons, bold *p*-values are significant. Asterisks indicate significance levels: *p*\* < 0.05, *p*\*\* < 0.01, *p*\*\*\* < 0.001.

After additionally accounting for the effect of log-transformed WMH volume, there was no effect of confluence in deep WM on cognitive performance, while for periventricular WM confluence, an effect was identified on all cognitive domains, and for whole-brain confluence an association was found with all subtests except for visuospatial function, working memory and the PACC score.

WMH confluence differed between all pairs of diagnostic groups except for SCD vs. MCI in the case of deep WMH.

## 5 Discussion

### 5.1 Overview

In this study, we present a novel and fully automated rater-independent metric for quantifying the confluence of white matter hyperintensities in MRI scans, a clinically relevant radiological marker which could previously only be assessed with visual rating scales. The confluence metric assigns each person a scalar value between 0 and 1 and is based on WMH segmentations. We investigated the relationship between WMH confluence and WMH volume, Fazekas scores, age and cognition, as well as differences in confluence between diagnostic groups, in QMIN-MC, a real world dataset of individuals attending UK memory clinics with a variety of neurodegenerative and non-degenerative cognitive disorders. We identified the following key findings: WMH confluence was highly correlated with WMH volume, Fazekas scores and age. Confluence is individually associated with performance across cognitive domains including memory, and is greater in vascular dementia, but does not explain cognitive performance in addition to WMH volume in this population. We validated these findings in an independent dataset from the German DELCODE study designed to investigate early signs of dementia.

### 5.2 In a population with neurodegenerative diseases, WMH confluence is highly correlated with WMH volume

Despite the confluence metric being independent of lesion volume, demonstrated with theoretical examples and exemplary images in Section 2.3, overall WMH confluence was found to be highly correlated with log-transformed WMH volume in the memory clinic cohorts investigated in this study. This indicates that as WMH grow in size, their topology shifts from punctate to increasingly confluent, which is consistent with how lesion volume tends to increase in progressive white matter injury: while new lesions do develop in previously unaffected sites, the predominant mechanism of lesion growth is an outward expansion of existing lesions. In this context, the term “WMH penumbra” was introduced by Maillard et al. (2011) to refer to the tissue surrounding WMH which displays more subtle pathology and is at risk of converting into visibly lesioned tissue (Voorter et al., 2025; Duering et al., 2013; Bergamino et al., 2025; Promjunyakul et al., 2018; Kern et al., 2023; Ferris et al., 2022). Penumbra tissue has been shown to decline in integrity over time (Maillard et al. (2014)), and Maillard et al. (2012) identified that 80% of incident WMH voxels constituted extensions of existing lesions. With respect to our study, this implies that an increase in volume will typically be associated with an increase in lesion confluence. The theoretically conceivable case of individuals with equal lesion volume and meaningfully different degrees of confluence was rare in the cohorts examined here. In other pathologies such as multiple sclerosis (MS), increases in lesion volume are driven by other mechanisms compared to neurodegenerative or cerebrovascular diseases. In MS, early disease (acute and relapsing MS) is characterised by the development of active lesions in new focal sites of neuroinflammation and demeyelination rather than due to outward expansion of existing lesions (Lassmann, 2019). This implies a weaker relationship between lesion volume and confluence, and makes the application of the confluence metric to an MS cohort an interesting avenue for future work.

### 5.3 WMH confluence is highly correlated with Fazekas scores and increases with age

The confluence metric’s validity was supported by demonstrating correlation with visual Fazekas scale ratings, suggesting that the metric accurately captures the concept of WMH confluence and severity. In line with previous studies reporting progression of WMH over time in diagnostic groups, we identified an increase of whole-brain WMH confluence with age (Jochems et al., 2022).

### 5.4 Higher whole-brain WMH confluence is associated with lower performance across cognitive domains

When considering confluence individually, we found a relationship of whole-brain WMH confluence with performance across cognitive domains, including attention, memory, fluency, language and visuospatial performance, as well as global cognition. This was the case in both the QMIN-MC and the DELCODE cohorts despite their differences in cognitive tests and represented diagnostic groups. This finding is consistent with our hypothesis that more confluent lesions contribute to more serious cognitive impairment because they disrupt a broader network of interconnected brain regions than punctate lesions. This mechanistic hypothesis can be explicitly explored in future work by relating our confluence metric to measures of white matter tract disruption and disconnectivity. However, due to the nature of the correlation between confluence and volume, it is difficult to disentangle the two effects.

Our findings are consistent with previous studies which have used other methods to characterise WMH. A few studies have focused on WMH confluence using visual assessment or by dichotomising existing visual rating scales. Kumar et al. (2022) found that among APOE4 carriers with MCI/SCI, individuals with confluent WMH had 7.85 greater odds of developing episodic memory impairment than individuals with non-confluent WMH. In MCI patients, Kumar et al. (2020) identified a link between confluent WMH and functional connectivity changes in fronto-parietal, fronto-temporal and temporo-parietal areas which could impair cognition. Heng et al. (2021) found that MCI patients with confluent WMH declined more rapidly in global cognition than those with non-confluent WMH and had greater risk of progressing to dementia.

In cognitively normal participants, Xing et al. (2022) found that confluent WMH affected memory and naming tasks more than mild WMH. In contrast, van Straaten et al. (2006) compared 46 individuals with punctate and confluent WMH of similar volumes and found no association with a range of symptoms including memory. Similarly, O’Brien et al. (1997) found no association between visually rated WMH and memory function. van Straaten et al. (2006) concluded that the lack of correlation with clinical impairment reflected the low granularity of visual rating scales and their inability to distinguish between individuals with different lesion loads. Our quantitative approach overcomes this limitation.

When considering whole-brain WMH confluence and log-transformed (but not raw) volume jointly, neither confluence nor volume showed an association with cognitive performance. Given collinearity between confluence and volume in this population, either of the two metrics individually is better suited to explain cognition. Confluence does not contain additional information regarding cognitive performance, however, as it is measured on a scale between 0 and 1, it might be more intuitive and easy to use as a metric compared to volume on a logarithmic scale.

Relatively low *R*^2^ values in the QMIN-MC cohort indicate that WMH volume and confluence explain relatively little variance in cognitive performance, in line with Frisoni et al. (2007) and Kloppenborg et al. (2014). Conversely, *R*^2^ values in the DELCODE group were larger. This discrepancy could be explained by the different composition of the two cohorts in terms of disease stage and severity of cognitive impairment. For the more cognitively impaired participants in QMIN-MC, other neurodegenerative processes might explain a higher proportion of cognitive performance than white matter damage can. On the other hand, the DELCODE cohort with its focus on the early stages of disease including subjective cognitive decline represents participants with less progressed neurodegenerative processes. Therefore, a larger proportion of cognitive impairment could be attributable to white matter changes.

### 5.5 Periventricular WMH are associated with performance in more cognitive domains than deep WMH

Considering evidence that deep and periventricular white matter lesions may represent separate pathological entities, we investigated deep and periventricular WMH confluence separately, finding the same relationships as whole-brain WMH confluence with respect to age and vascular dementia vs. other diagnoses. With respect to the relationship with cognition, in the QMIN-MC cohort we found an association of periventricular WMH confluence with performance in all cognitive domains, while deep WMH confluence was associated with memory, fluency and language performance. Similarly, in the DELCODE cohort we identified a relationship of periventricular WMH confluence across cognitive domains, while deep WMH confluence had no relationship with any domain. When considering lesion volume in addition to confluence, only an effect of periventricular lesion volume on fluency and language performance remained.

A number of previous studies have investigated deep and periventricular WMH and their role in cognitive impairment separately. Most studies have identified effects in PWMH only (J. C. De Groot et al., 2002, Debette et al., 2007; Heuvel et al., 2006;Prins et al., 2005, H. J. Yang et al., 2024,Melazzini et al., 2021, Kim et al., 2011, Gronewold et al., 2022,Huang et al., 2018), while a small number of studies have reported effects for DWMH only (Soriano-Raya et al., 2012, Yoon et al., 2018), or no difference between PWMH and DWMH in the relationships with cognition (Garde et al., 2000, Burns et al., 2005, Courtney et al., 2025). As mentioned by Vergoossen et al. (2020) and J. De Groot et al. (2000), the stronger involvement of periventricular WMH as opposed to deep WMH in cognitive impairment could be due to the fact that the location of PWMH overlaps with multiple long association tracts which support cognitive functions, while DWMH due to their location would disrupt short cortico-cortical tracts and local networks. Indeed the locations of PWMH are typically traversed by the long arcuate fasciculus, superior and inferior longitudinal fasciculus, inferior fronto-occipital fasciculus and the cingulum.

It should be noted that multiple different criteria have been used for separating white matter into deep and periventricular regions, e.g. various distances from ventricle edges between 3 and 13 mm, continuity from the lateral ventricles or separation with unsupervised algorithms. There is no consensus in the literature as to whether and how WMH should be analysed based on their location (Herńandez et al., 2014; Valdés Herńandez et al., 2012), and some studies have argued that such a separation is arbitrary and does not relate to subpopulations of WMH (DeCarli et al., 2005; Barkhof and Scheltens, 2006), while others have recommended considering the two regions separately (Garde et al., 2000). Therefore it is unclear whether heterogeneous findings reflect methodological differences or genuine biological differences in the effects of PWMH and DWMH.

### 5.6 WMH confluence is higher in vascular dementia than in other diagnostic groups, and lowest in participants with no cognitive impairment

Confluence scores did not add to the limited ability of WMH lesion volume to distinguish between diagnostic groups. Investigating different diagnostic groups in the QMIN-MC cohort, we found that participants with vascular or mixed dementia and with no cognitive impairment differed in WMH confluence scores from those with all other diagnoses. Similar results were obtained for WMH volume. Given that white matter lesions are the hallmark of vascular dementia and a diagnostic criterion, this finding is not surprising and in line with previous work (Luo et al., 2025). However, beyond this there is no clear role for WMH confluence in distinguishing between diagnostic groups. Confluence scores cover a wide range within diagnostic groups and show large overlap between groups, suggesting that confluence is associated with severity across clinical populations rather than being a disease-specific pathology.

### 5.7 Strengths of the confluence metric

The proposed confluence metric has various strengths over currently used visual rating scales. Firstly, since it is fully automated and can be calculated without user input, it is reproducible, objective and reliable. Currently available visual rating scales routinely used in research and clinical practice, such as the Fazekas scale (Fazekas et al., 1993), the Scheltens scale (Scheltens et al., 1993) or the ARWMS scale (Wahlund et al., 2001), are subjective, time-consuming, and suffer from low inter/intra-rater reliability (van den Heuvel et al., 2006; Prins et al., 2004; Kapeller et al., 2003). A few previous studies investigating the effect of confluent WMH have dichotomised the scores obtained with visual rating scales using arbitrary cut-offs in order to provide classifications of “punctate WMH” and “confluent WMH”. Such classifications are at risk of low transparency and low reliability.

Secondly, the confluence metric is continuous and offers a fine-grained distinction of WMH, particularly at the more severe end. This can be seen in Figure 4 where QMIN-MC participants in the Fazekas 3 group demonstrate a clear ceiling effect with a wide distribution of confluence scores. Similarly, Fazekas group 1 in the DELCODE group is composed of participants with a very wide range of confluence scores who, despite differences in phenotype and cognitive performance, all receive the same score on a visual rating scale. This may explain why visual scales are not as sensitive for capturing WMH progression (Enzinger et al., 2007), distinguishing between diagnostic groups (Enzinger et al., 2006) or identifying associations with clinical features (van Straaten et al., 2006).

### 5.8 Strengths and limitations of the study

A key strength of this study is the validation in an independent dataset including participants with different characteristics, a different MRI scanner and protocol, and processed using a different WMH segmentation algorithm. We show the versatility of the tool given that no MRI scans were excluded because of data quality. In addition, we demonstrate WMH confluence is potentially relevant across a wide age range and a mixed range of neurodegenerative conditions, non-progressive cognitive disorders, and cognitively normal elderly individuals.

It is a limitation of the study that due to the cross-sectional nature of the dataset no causal inference can be made as to the effect of confluence on cognition. We could not evaluate the impact of confluence on disease progression or future risk. This poses an interesting opportunity for future work with longitudinal datasets.

### 5.9 Concluding remarks

We present a novel tool to quantify confluence of white matter hyperintensities relevant to age and cognition. WMH confluence individually was associated with cognitive performance across domains, but was highly correlated with WMH volume in this population and did therefore not explain any variance in addition to volume. In participants with vascular dementia, confluence was higher than in those with other neurodegenerative diseases, while in participants with no cognitive impairment it was lower than in the cognitively impaired. Confluence scores are strongly correlated with Fazekas ratings, but contain additional information and granularity. Therefore our confluence metric offers a reproducible and quantitative metric that can complement visual rating scales in studies investigating white matter disease.

## Data and Code Availability

The imaging and clinical data drawn from QMIN-MC is available on DPUK. Original Python code is available on GitHub here: https://github.com/MRTanja/WMH confluence.

## Author Contributions

T.S.: Conceptualisation, Methodology, Code, Formal analysis, Data curation (QMIN-MC), Writing—Original Draft. R.S.: Methodology, Writing—Review and Editing. M.M.: Data curation (QMIN-MC), Writing—Review and Editing. D.C.: Study design, Writing—Review and Editing. J.B.: Formal analysis, Data curation (DELCODE), Writing-Review and Editing. M.P.: Data curation (DELCODE). P.A.: Data curation (DELCODE). O.P.: Data curation (DELCODE). J.H.R.: Data curation (DELCODE). L.P.: Data curation (DELCODE). D.G.: Data curation (DELCODE). J.P.: Data curation (DELCODE). E.J.S.: Data curation (DELCODE). M.G.: Data curation (DELCODE). S.A.: Data curation (DELCODE). A.S.: Data curation (DELCODE). K.F.: Data curation (DELCODE). O.K.: Data curation (DELCODE). J.W.: Data curation (DELCODE). C.B.: Data curation (DELCODE). B.H.S.: Data curation (DELCODE). A.R.: Data curation (DELCODE). W.G.: Data curation (DELCODE). E.I.I: Data curation (DELCODE). M.B.: Data curation (DELCODE). K.B.: Data curation (DELCODE). D.J.: Data curation (DELCODE). S.S.: Data curation (DELCODE). R.P.: Data curation (DELCODE). B.S.R.: Data curation (DELCODE). S.T.: Data curation (DELCODE). M.M.: Data curation (DELCODE). A.G.: Data curation (DELCODE). C.L.: Data curation (DELCODE). S.S.: Data curation (DELCODE). A.S.: Data curation (DELCODE). G.C.P.: Data curation (DELCODE). M.W.: Data curation (DELCODE). F.L.: Data curation (DELCODE). L.K.: Data curation (DELCODE). S.H.: Data curation (DELCODE). P.D.: Data curation (DELCODE). S.S.: Data curation (DELCODE). E.D.: Data curation (DELCODE). F.J.: Data curation (DELCODE). G.Z.: Data curation (DELCODE). B.R.U.: Study design, Writing—Review and Editing. T.R.: Conceptualisation, Writing—Review and Editing, Supervision.

## Funding

This research was supported by Gates Cambridge, Alzheimer’s Research UK (ARUK-SRF2023B-005), and the National Institute for Health and Care Research (NIHR) Cambridge Biomedical Research Centre (NIHR203312). The views expressed are those of the authors and not necessarily those of the NIHR or the Department of Health and Social Care.

## Declaration of Competing Interests

E.D. reports personal fees from Biogen, Roche, Lilly, Eisai and UCL Consultancy, as well as non-financial support from Rox Health. He is scientific co-founder of neotiv GmbH and owns company shares. C.B. received honoraria as a commercial advisory board member for Lilly and honoraria for lectures from Boehringer Ingelheim, Novo Nordisk, Lilly and Eisai. All other authors declare no non-financial or financial competing interests.

## Acknowledgements

T.S. would like to extend a special thank you to Filippo De Luca for his helpful mathematical insights, and to William Underwood for the valuable statistical advice.

## Footnotes

1 More recently shape metrics such as concavity and eccentricity have been proposed to further characterise the morphological features of WMH (de Bresser et al., 2018; Ghaznawi et al., 2019; Keller et al., 2022; Lange et al., 2017; Gwo et al., 2019; Kant et al., 2019). These will not be focused on here.

## Notes

### Competing Interest Statement

The authors have declared no competing interest.

