## Supplemental Materials for "Automated quantification of white matter hyperintensity confluence: A measure of spatial organisation beyond volume and visual rating scales"

### 1 Supplementary Materials

Table 1: Relationship between ACE subtests and WMH: Regression Model 1 with confluence as a predictor. QMIN-MC data[a].

| | | Whole-brain WMH confluence | | | | | DWMH confluence | | | | | PWMH confluence | | | | |
|---|---|---|---|---|---|---|---|---|---|---|---|---|---|---|---|---|
| Outcome variable | Predictor | $\beta$ | SD | $t$ | $p_{FDR}$ | $R^2$ | $\beta$ | SD | $t$ | $p_{FDR}$ | $R^2$ | $\beta$ | SD | $t$ | $p_{FDR}$ | $R^2$ |
| Total ACE | Intercept | 77.81 | 7.45 | 10.44 | <**0.001***** | 0.09 | 89.79 | 7.06 | 12.72 | <**0.001***** | 0.07 | 77.35 | 7.43 | 10.41 | <**0.001***** | 0.10 |
| | Age | -0.19 | 0.06 | -3.04 | **0.005**** | | -0.32 | 0.06 | -5.63 | <**0.001***** | | -0.18 | 0.06 | -2.86 | **0.008**** | |
| | Sex (male) | 2.25 | 1.34 | 1.68 | 0.124 | | 2.66 | 1.36 | 1.95 | 0.074 | | 2.29 | 1.34 | 1.71 | 0.120 | |
| | Confluence | -35.07 | 6.71 | -5.23 | <**0.001***** | | -25.44 | 9.71 | -2.62 | **0.016*** | | -31.68 | 5.83 | -5.43 | <**0.001***** | |
| | Intracranial volume | 0.00 | 0.00 | 1.51 | 0.167 | | 0.00 | 0.00 | 1.02 | 0.361 | | 0.00 | 0.00 | 1.46 | 0.177 | |
| Attention | Intercept | 22.07 | 1.35 | 16.39 | <**0.001***** | | 24.24 | 1.24 | 19.58 | <**0.001***** | | 21.95 | 1.35 | 16.22 | <**0.001***** | |
| | Age | -0.09 | 0.02 | -5.11 | <**0.001***** | | -0.12 | 0.02 | -7.45 | <**0.001***** | | -0.09 | 0.02 | -5.00 | <**0.001***** | |
| | Sex (male) | 0.84 | 0.38 | 2.23 | **0.039*** | | 0.96 | 0.38 | 2.52 | **0.020*** | | 0.85 | 0.38 | 2.26 | **0.037*** | |
| | Confluence | -6.86 | 1.86 | -3.69 | <**0.001***** | | -1.97 | 2.67 | -0.74 | 0.511 | | -6.11 | 1.62 | -3.77 | <**0.001***** | |
| | Intracranial volume | 0.44 | 0.19 | 2.31 | **0.034*** | | 0.35 | 0.19 | 1.85 | 0.092 | | 0.43 | 0.19 | 2.26 | **0.037*** | |
| Memory | Intercept | 18.18 | 2.80 | 6.49 | <**0.001***** | 0.08 | 21.72 | 2.64 | 8.23 | <**0.001***** | 0.06 | 18.25 | 2.79 | 6.53 | <**0.001***** | 0.08 |
| | Age | -0.07 | 0.02 | -3.06 | **0.005**** | | -0.11 | 0.02 | -5.25 | <**0.001***** | | -0.07 | 0.02 | -3.00 | **0.005**** | |
| | Sex (male) | 0.77 | 0.50 | 1.53 | 0.163 | | 0.87 | 0.51 | 1.71 | 0.120 | | 0.79 | 0.50 | 1.58 | 0.151 | |
| | Confluence | -11.78 | 2.53 | -4.66 | <**0.001***** | | -11.00 | 3.63 | -3.03 | **0.005**** | | -10.21 | 2.20 | -4.64 | <**0.001***** | |
| | Intracranial volume | 0.00 | 0.00 | 0.94 | 0.397 | | 0.00 | 0.00 | 0.60 | 0.594 | | 0.00 | 0.00 | 0.87 | 0.434 | |
| Fluency | Intercept | 44.94 | 3.98 | 11.30 | <**0.001***** | 0.11 | 50.33 | 3.75 | 13.42 | <**0.001***** | 0.09 | 44.77 | 3.96 | 11.29 | <**0.001***** | 0.11 |
| | Age | -0.17 | 0.03 | -4.88 | <**0.001***** | | -0.22 | 0.03 | -7.35 | <**0.001***** | | -0.16 | 0.03 | -4.73 | <**0.001***** | |
| | Sex (male) | 1.05 | 0.72 | 1.45 | 0.178 | | 1.23 | 0.73 | 1.69 | 0.123 | | 1.07 | 0.72 | 1.48 | 0.173 | |
| | Confluence | -16.33 | 3.60 | -4.54 | <**0.001***** | | -12.64 | 5.16 | -2.45 | **0.024*** | | -14.67 | 3.13 | -4.69 | <**0.001***** | |
| | Intracranial volume | 0.00 | 0.00 | 0.05 | 0.970 | | -0.00 | 0.00 | -0.35 | 0.762 | | 0.00 | 0.00 | 0.00 | 0.999 | |
| Language | Intercept | 21.03 | 1.47 | 14.27 | <**0.001***** | | 23.45 | 1.34 | 17.56 | <**0.001***** | | 20.76 | 1.48 | 14.03 | <**0.001***** | |
| | Age | 0.04 | 0.02 | 1.84 | 0.092 | | 0.00 | 0.02 | 0.19 | 0.865 | | 0.04 | 0.02 | 2.02 | 0.064 | |
| | Sex (male) | 0.21 | 0.43 | 0.49 | 0.661 | | 0.29 | 0.43 | 0.68 | 0.545 | | 0.22 | 0.43 | 0.52 | 0.646 | |
| | Confluence | -9.66 | 2.13 | -4.54 | <**0.001***** | | -8.29 | 3.03 | -2.74 | **0.011*** | | -8.88 | 1.85 | -4.80 | <**0.001***** | |
| | Intracranial volume | 0.70 | 0.21 | 3.25 | **0.003**** | | 0.62 | 0.22 | 2.91 | **0.007**** | | 0.68 | 0.21 | 3.21 | **0.003**** | |
| Visuospatial | Intercept | 17.02 | 1.22 | 13.97 | <**0.001***** | | 18.40 | 1.11 | 16.57 | <**0.001***** | | 16.75 | 1.22 | 13.69 | <**0.001***** | |
| | Age | -0.04 | 0.02 | -2.58 | **0.017*** | | -0.06 | 0.01 | -4.15 | <**0.001***** | | -0.04 | 0.02 | -2.34 | **0.032*** | |
| | Sex (male) | 1.64 | 0.34 | 4.76 | <**0.001***** | | 1.73 | 0.35 | 5.01 | <**0.001***** | | 1.63 | 0.34 | 4.75 | <**0.001***** | |
| | Confluence | -3.77 | 1.71 | -2.21 | **0.041*** | | 0.51 | 2.45 | 0.21 | 0.863 | | -3.86 | 1.48 | -2.61 | **0.016*** | |
| | Intracranial volume | 0.22 | 0.17 | 1.30 | 0.231 | | 0.16 | 0.17 | 0.94 | 0.397 | | 0.22 | 0.17 | 1.31 | 0.227 | |

[a] Predictors included in the model: WMH confluence, age, sex, total intracranial volume. *P*-values have been FDR-corrected for 90 tests, bold *p*-values are significant. Asterisks indicate significance levels: p* $<$ 0.05, p** $<$ 0.01, p*** $<$ 0.001. No $R^2$ is available for Tobit regressions (attention, language and visuospatial abilities).

Table 2: Relationship between ACE subtests and WMH: Regression model 2 with confluence and volume as predictors. QMIN-MC data[a].

| | | Whole-brain WMH Confluence | | | | | DWMH Confluence | | | | | PWMH Confluence | | | | |
|---|---|---|---|---|---|---|---|---|---|---|---|---|---|---|---|---|
| Outcome variable | Predictor | $\beta$ | SD | $t$ | $p_{FDR}$ | $R^2$ | $\beta$ | SD | $t$ | $p_{FDR}$ | $R^2$ | $\beta$ | SD | $t$ | $p_{FDR}$ | $R^2$ |
| Total ACE | Intercept | 63.70 | 7.89 | 8.07 | **<0.001*** ** | 0.12 | 82.35 | 7.31 | 11.26 | **<0.001*** ** | 0.08 | 64.92 | 7.88 | 8.24 | **<0.001*** ** | 0.12 |
| | Age | -0.06 | 0.07 | -0.88 | 0.462 | | -0.25 | 0.06 | -4.04 | **<0.001*** ** | | -0.07 | 0.07 | -0.97 | 0.429 | |
| | Sex (male) | 1.78 | 1.33 | 1.34 | 0.272 | | 2.43 | 1.35 | 1.80 | 0.145 | | 2.04 | 1.33 | 1.54 | 0.210 | |
| | Volume | -2.75 | 1.35 | -2.04 | 0.093 | | -1.94 | 0.71 | -2.72 | **0.022*** | | -3.04 | 1.69 | -1.80 | 0.145 | |
| | Confluence | -16.83 | 18.77 | -0.90 | 0.462 | | 5.88 | 20.88 | 0.28 | 0.811 | | -8.24 | 20.50 | -0.40 | 0.741 | |
| | Volume*Confluence | 13.13 | 5.16 | 2.54 | **0.031*** | | 4.39 | 4.93 | 0.89 | 0.462 | | 8.34 | 5.35 | 1.56 | 0.206 | |
| | Intracranial volume | 0.00 | 0.00 | 2.08 | 0.087 | | 0.00 | 0.00 | 1.34 | 0.272 | | 0.00 | 0.00 | 1.96 | 0.110 | |
| Attention | Intercept | 18.83 | 1.46 | 12.86 | **<0.001*** ** | | 22.73 | 1.30 | 17.53 | **<0.001*** ** | | 19.21 | 1.46 | 13.14 | **<0.001*** ** | |
| | Age | -0.06 | 0.02 | -2.89 | **0.013*** | | -0.10 | 0.02 | -5.95 | **<0.001*** ** | | -0.06 | 0.02 | -3.04 | **0.008** ** | |
| | Sex (male) | 0.72 | 0.37 | 1.94 | 0.112 | | 0.91 | 0.38 | 2.43 | **0.040*** | | 0.79 | 0.37 | 2.14 | 0.080 | |
| | Volume | -0.40 | 0.38 | -1.05 | 0.392 | | -0.49 | 0.20 | -2.44 | **0.039*** | | -0.67 | 0.47 | -1.41 | 0.255 | |
| | Confluence | -7.51 | 5.19 | -1.45 | 0.246 | | 5.14 | 5.98 | 0.86 | 0.469 | | -2.11 | 5.70 | -0.37 | 0.754 | |
| | Volume*Confluence | 5.38 | 1.47 | 3.67 | **<0.001*** ** | | 1.62 | 1.41 | 1.15 | 0.353 | | 3.13 | 1.51 | 2.08 | 0.087 | |
| | Intracranial volume | 0.51 | 0.19 | 2.70 | **0.022*** | | 0.40 | 0.19 | 2.09 | 0.087 | | 0.50 | 0.19 | 2.60 | **0.027*** | |
| Memory | Intercept | 12.82 | 2.96 | 4.33 | **<0.001*** ** | 0.11 | 19.59 | 2.74 | 7.14 | **<0.001*** ** | 0.07 | 13.80 | 2.95 | 4.67 | **<0.001*** ** | 0.11 |
| | Age | -0.02 | 0.03 | -0.88 | 0.462 | | -0.09 | 0.02 | -3.99 | **<0.001*** ** | | -0.03 | 0.03 | -1.13 | 0.358 | |
| | Sex (male) | 0.58 | 0.50 | 1.17 | 0.352 | | 0.80 | 0.51 | 1.58 | 0.202 | | 0.69 | 0.50 | 1.39 | 0.261 | |
| | Volume | -0.62 | 0.51 | -1.23 | 0.324 | | -0.51 | 0.27 | -1.91 | 0.120 | | -0.28 | 0.63 | -0.44 | 0.720 | |
| | Confluence | -10.89 | 7.04 | -1.55 | 0.208 | | -3.90 | 7.83 | -0.50 | 0.690 | | -12.20 | 7.69 | -1.59 | 0.202 | |
| | Volume*Confluence | 6.76 | 1.94 | 3.48 | **0.002** ** | | 1.91 | 1.85 | 1.03 | 0.400 | | 6.33 | 2.01 | 3.14 | **0.006** ** | |
| | Intracranial volume | 0.00 | 0.00 | 1.42 | 0.255 | | 0.00 | 0.00 | 0.84 | 0.475 | | 0.00 | 0.00 | 1.08 | 0.379 | |
| Fluency | Intercept | 37.54 | 4.21 | 8.92 | **<0.001*** ** | 0.13 | 46.14 | 3.88 | 11.89 | **<0.001*** ** | 0.11 | 38.02 | 4.20 | 9.05 | **<0.001*** ** | 0.13 |
| | Age | -0.10 | 0.04 | -2.61 | **0.027*** | | -0.18 | 0.03 | -5.50 | **<0.001*** ** | | -0.10 | 0.04 | -2.69 | **0.022*** | |
| | Sex (male) | 0.84 | 0.71 | 1.18 | 0.347 | | 1.13 | 0.72 | 1.56 | 0.206 | | 0.96 | 0.71 | 1.36 | 0.270 | |
| | Volume | -1.96 | 0.72 | -2.71 | **0.022*** | | -1.24 | 0.38 | -3.28 | **0.004** ** | | -2.11 | 0.90 | -2.34 | **0.049*** | |
| | Confluence | 0.72 | 10.02 | 0.07 | 0.950 | | 11.01 | 11.10 | 0.99 | 0.422 | | 4.04 | 10.93 | 0.37 | 0.754 | |
| | Volume*Confluence | 4.86 | 2.76 | 1.76 | 0.155 | | 0.43 | 2.62 | 0.16 | 0.891 | | 2.74 | 2.87 | 0.96 | 0.437 | |
| | Intracranial volume | 0.00 | 0.00 | 0.67 | 0.584 | | -0.00 | 0.00 | -0.02 | 0.980 | | 0.00 | 0.00 | 0.61 | 0.614 | |
| Language | Intercept | 18.15 | 1.63 | 11.12 | **<0.001*** ** | | 21.85 | 1.42 | 15.42 | **<0.001*** ** | | 18.17 | 1.63 | 11.16 | **<0.001*** ** | |
| | Age | 0.07 | 0.02 | 3.29 | **0.004** ** | | 0.03 | 0.02 | 1.40 | 0.259 | | 0.07 | 0.02 | 3.15 | **0.006** ** | |
| | Sex (male) | 0.11 | 0.42 | 0.26 | 0.823 | | 0.26 | 0.43 | 0.61 | 0.614 | | 0.17 | 0.42 | 0.40 | 0.741 | |
| | Volume | -0.70 | 0.43 | -1.63 | 0.192 | | -0.72 | 0.23 | -3.14 | **0.006** ** | | -0.17 | 0.54 | -0.32 | 0.784 | |
| | Confluence | -5.30 | 5.96 | -0.89 | 0.462 | | 7.39 | 6.69 | 1.10 | 0.369 | | -10.42 | 6.51 | -1.60 | 0.200 | |
| | Volume*Confluence | 3.50 | 1.65 | 2.12 | 0.081 | | -1.13 | 1.56 | -0.72 | 0.552 | | 4.13 | 1.71 | 2.42 | **0.040*** | |
| | Intracranial volume | 0.78 | 0.21 | 3.64 | **<0.001*** ** | | 0.67 | 0.21 | 3.14 | **0.006** ** | | 0.71 | 0.22 | 3.27 | **0.004** ** | |
| Visuospatial | Intercept | 14.38 | 1.33 | 10.84 | **<0.001*** ** | | 17.20 | 1.17 | 14.66 | **<0.001*** ** | | 14.42 | 1.32 | 10.93 | **<0.001*** ** | |
| | Age | -0.01 | 0.02 | -0.63 | 0.611 | | -0.05 | 0.02 | -2.92 | **0.012*** | | -0.01 | 0.02 | -0.58 | 0.635 | |
| | Sex (male) | 1.54 | 0.34 | 4.53 | **<0.001*** ** | | 1.69 | 0.34 | 4.92 | **<0.001*** ** | | 1.58 | 0.34 | 4.66 | **<0.001*** ** | |
| | Volume | -0.58 | 0.34 | -1.68 | 0.176 | | -0.38 | 0.18 | -2.06 | 0.088 | | -0.81 | 0.43 | -1.87 | 0.127 | |
| | Confluence | -0.60 | 4.77 | -0.13 | 0.915 | | 6.21 | 5.35 | 1.16 | 0.352 | | 2.57 | 5.24 | 0.49 | 0.690 | |
| | Volume*Confluence | 3.44 | 1.33 | 2.58 | **0.028*** | | 1.10 | 1.26 | 0.87 | 0.463 | | 1.87 | 1.38 | 1.35 | 0.270 | |
| | Intracranial volume | 0.29 | 0.17 | 1.72 | 0.163 | | 0.20 | 0.17 | 1.15 | 0.353 | | 0.30 | 0.17 | 1.75 | 0.157 | |

[a] Predictors included in the model: WMH confluence, WMH volume, WMH volume × WMH confluence, age, sex, total intracranial volume. $P$-values have been FDR-corrected for 126 tests, bold $p$-values are significant. Asterisks indicate significance levels: p* $< 0.05$, p** $< 0.01$, p*** $< 0.001$. No $R^2$ is available for Tobit regressions (attention, language and visuospatial abilities).
